# Reconstructing human Brown Fat developmental trajectory in vitro

**DOI:** 10.1101/2022.06.01.494355

**Authors:** Jyoti Rao, Jerome Chal, Fabio Marchianò, Chih-Hao Wang, Ziad Al Tanoury, Svetlana Gapon, Yannis Djeffal, Alicia Mayeuf-Louchart, Ian Glass, Elizabeth M. Sefton, Bianca Habermann, Gabrielle Kardon, Fiona M. Watt, Yu-Hua Tseng, Olivier Pourquié

## Abstract

Brown adipocytes represent a specialized type of mammalian adipocytes able to uncouple nutrient catabolism from ATP generation to dissipate energy as heat. They play an important role in mammals, allowing non-shivering thermogenesis to regulate body temperature in response to cold exposure. In humans, the brown fat tissue is composed of small discrete depots found mostly throughout the neck and trunk region. Increasing brown fat activity either with drug treatment or cell therapy is considered a potential approach for the treatment of metabolic syndrome and obesity. The recent development of in vitro differentiation strategies relying on human pluripotent stem cells (hPSCs) offers the possibility to produce unlimited amounts of brown adipocytes. A strategy efficiently applied to several tissues is to recapitulate step by step the development of the tissue of interest by exposing hPSCs to the signaling cues used during normal embryonic development. However, this strategy has proven difficult to implement for brown fat as the development of this tissue is poorly understood. Here, we first used single cell RNA sequencing to characterize the development of interscapular brown fat in mouse. Our analysis identified a previously unrecognized population of brown adipocytes precursors characterized by expression of the transcription factor GATA6. We show that this precursor population can be efficiently generated from paraxial mesoderm precursors differentiated in vitro from hPSCs by modulating the signaling pathways identified in our transcriptomic analysis. These precursors can in turn be efficiently converted into functional brown adipocytes which can respond to adrenergic stimuli by increasing their metabolism resulting in heat production.

## INTRODUCTION

In humans, brown adipose tissue (BAT) is present most abundantly in the interscapular region at birth and in the supraclavicular region in adulthood (Heaton, 1972; Lidell, 2019; Merklin, 1974; van Marken Lichtenbelt et al., 2009; Virtanen et al., 2009). Brown adipocytes derive from the paraxial mesoderm, an embryonic tissue that flanks the neural tube. Its main derivatives are the axial skeleton, the skeletal muscles and the dermis of the back. In mammals, the paraxial mesoderm first forms through gastrulation in the primitive streak and generates the presomitic mesoderm (PSM) in the posterior region of the embryo. As it matures, the PSM forms epithelial somites which become organized in two parallel arrays of segments providing the blueprint for the future metameric organization of vertebrae and other trunk derivatives. Soon after their formation, the ventral part of somites forms the mesenchymal sclerotome which yields the axial skeleton. The dorsal part of the somite remains epithelial, forming the dermomyotome. This compartment is composed of cells expressing the transcription factor Pax3, and it generates the dermis of the back and skeletal muscles of the body.

In mammals, the dermomyotome is also the main source of the brown adipocyte lineage. Lineage tracing experiments have shown that murine interscapular brown fat arises from Pax3, Meox1 and Myf5-expressing dermomyotomal cells in the embryo (Sanchez-Gurmaches and Guertin, 2014; Seale et al., 2008; Sebo et al., 2018; Wang et al., 2014). During development, Pax3-positive dermomyotome cells give rise to multipotential progenitor cells expressing Engrailed1 and Pax7. These multipotent cells develop into dermal cells, skeletal muscle and brown adipocytes (Atit et al., 2006; Lepper and Fan, 2010). Lineage tracing of Pax7 multipotent progenitors in mouse shows that these cells give rise to skeletal muscle, dermis and some brown adipocytes between E (embryonic day) 9.5 and E11.5 (Lepper and Fan, 2010; Sebo et al., 2018). The contribution of Pax7 progenitors becomes restricted to the skeletal muscle lineage at E11.5-12.5, suggesting that the brown fat lineage diverges from the skeletal muscle lineage around E11.5. Deletion of Pax7 in Pax7-expressing progenitors at E10.5 leads to increased contribution of these progenitors to BAT arguing for a repressive role of Pax7 on the brown adipocyte fate (An et al., 2017). Furthermore, determination of the adipogenic and myogenic fate involves mutual repression of Myod1/Myf5 and Prdm16 (An et al., 2017; Seale et al., 2008; Seale et al., 2007). Deletion of Myf5/Myod1 and Myog leads to an expansion of dorsal adipose depot (Hasty et al., 1993; Kablar et al., 2003). In contrast to skeletal muscle, it has been difficult to identify specific markers of brown fat precursors during development. Several genes such as Ebf2, Pdgfra, and Cd34 are expressed by adipocytes precursors but they are not specific for this cell population (Wang et al., 2014; Wang and Seale, 2016). Thus, the hierarchy of somitic precursors giving rise to this tissue is still poorly characterized. Overall, the development of the brown fat lineage in mouse remains poorly understood (Schulz and Tseng, 2013).

Our knowledge of human BAT development is even more limited. Current in vitro models to study human BAT are based on brown adipocytes differentiated in vitro from primary cell lines and stromal vascular cells (Samuelson and Vidal-Puig, 2020). Alternatively, several protocols have been established to differentiate human pluripotent stem cells (hPSCs) into brown adipocytes using either transgenic overexpression of transcription factors (Ahfeldt et al., 2012) or treatment with growth factors and small molecules (Takeda et al., 2017). Other reported methods rely on serum-based spontaneous differentiation or treatment of hPSCs with a cytokine cocktail (Guenantin et al., 2017; Hafner et al., 2016; Nishio et al., 2012). Recently, protocols allowing the generation of brown adipocytes from hPSCs by recapitulating developmental cues have also been reported (Carobbio et al., 2021; Zhang et al., 2020). However, to date, a well characterized roadmap of brown fat lineage development is still missing. Therefore, benchmarking these protocols against the normal trajectory of brown adipocytes differentiation in vivo has not been possible. In this study, we used single cell RNA sequencing (scRNAseq) to profile mouse somite-derived dorsal tissues (including brown fat, dermis and skeletal muscle) to investigate brown fat development. We identify a novel critical stage in brown adipocyte development characterized by the selective expression of the transcription factor Gata6. We previously developed an efficient protocol recapitulating the development of somitic mesoderm and skeletal muscle progenitors from human and mouse pluripotent stem cells in vitro (Chal et al., 2016; Chal et al., 2015; Diaz-Cuadros et al., 2020). Building on this protocol and using developmental cues identified in our scRNAseq data, we now report an efficient strategy to generate GATA6-positive brown adipocyte precursors and functional mature brown adipocytes in vitro from human induced pluripotent stem cells (iPSCs).

## RESULTS

### Single-cell transcriptomic analysis of the dorsal trunk of mouse embryos captures the development of somitic lineages

In mouse, interscapular brown adipose tissue (BAT) develops in between the dermis, trapezius muscle, pectoralis muscle, and deep dorsal muscle bundles. We first set out to analyze mouse interscapular brown fat development using immuno-histochemistry. We analyzed sections of the dorsal region of the developing mouse brachial trunk with markers of the adipocyte lineage. Developing adipocyte precursors expressing Pparg start to appear at E13.5 and develop in between the dermis and Myosin Heavy Chain (MyHC) expressing muscle fibers (Figure 1A). Perilipin1 (Plin1) staining shows that a small number of preadipocytes start to accumulate lipid vesicles at E14.5. An increasing number of lipid-containing immature adipocytes appears by E15.5 (Figure 1A).

**Figure 1.**
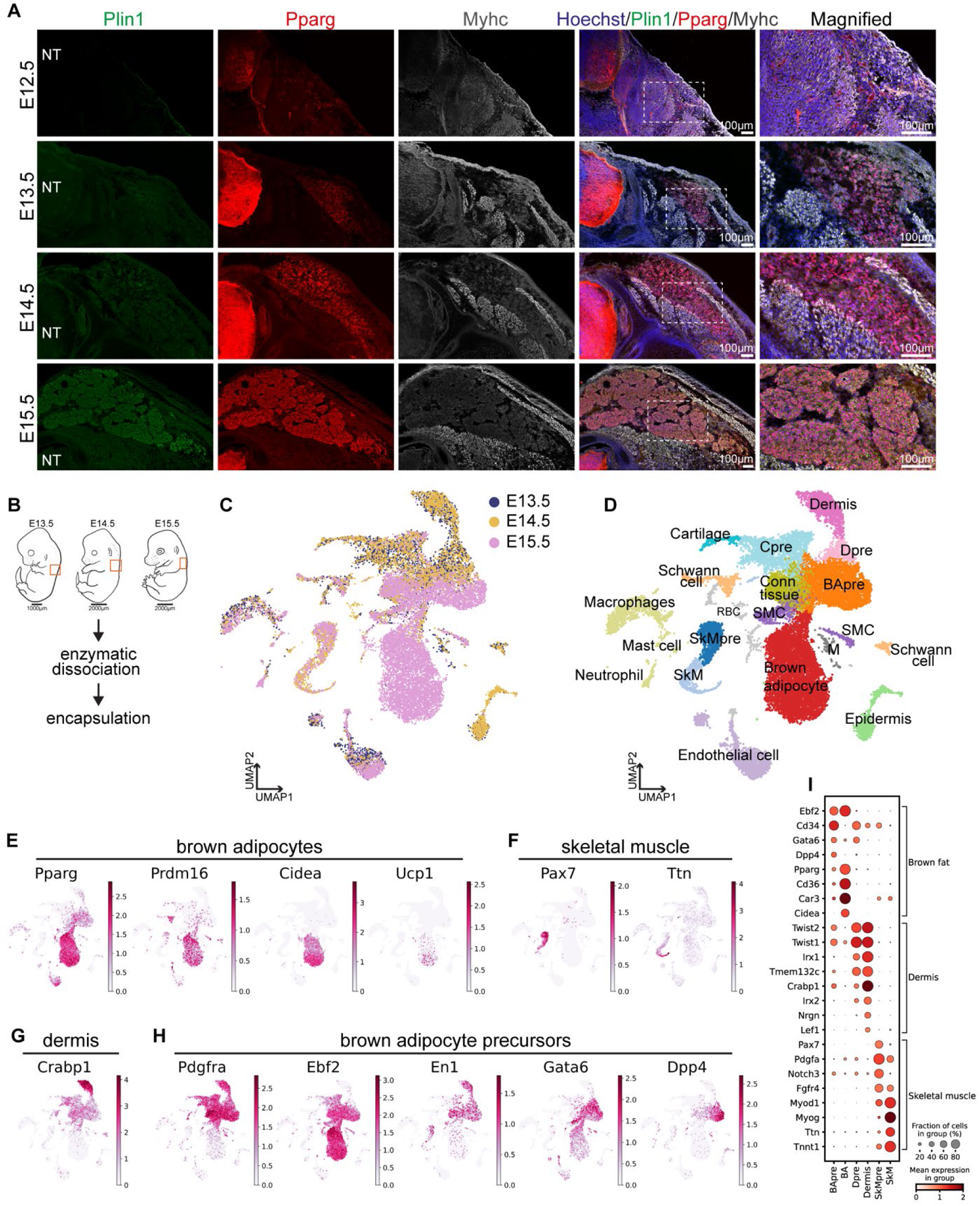
**A** – Immunofluorescence analysis of developing interscapular brown fat in mouse. Images represents transverse section of mouse embryos at the forelimb level showing interscapular region. Perilipin (Plin1) expressing adipocytes start to appear on embryonic day (E) 14.5 and perilipin is widely expressed in interscapular fat on E15.5. Staining for Pparg antibody illustrating expression of Pparg starting at E13.5. Interscapular fat develops in between muscle bundles stained using a Myosin heavy chain (Myhc) antibody. NT = Neural tube. **B** – Schematic representation of cell isolation strategy for single cell RNA sequencing. Dorsal interscapular region was dissected out with intact dermis and epidermis for embryonic day (E) 13.5 and 14.5. For E15.5, skin and underlying dermis was removed before cell dissociation. For each stage, tissues were isolated from two embryos. Tissues were digested with enzymes followed by encapsulation using inDrops platform. **C** – UMAP embedding of the single cells isolated from E13.5, E14.5 and E15.5 mouse embryos after Bkknn batch correction (50 PC dimensions, 20,000 cells) Colors indicate embryonic days. **D** – UMAP showing cell clusters identified using Leiden based clustering. Colors indicate identified cell cluster. BApre=brown adipocyte precursors, Conn tissue=connective tissue cells, SkM=skeletal muscle, SkMpre=skeletal muscle precursors, SMC=smooth muscle cells, Dpre= dermis precursors, Cpre=cartilage precursors, M=Meninges, RBC=red blood cells. **E** – Expression patterns of selected genes detected in the brown adipocyte cluster (BA). UMAP plots are colored by log-normalized transcript counts. **F** – Feature plots showing expression patterns of selected genes expressed in skeletal muscle precursor (SkMpre) and skeletal muscle cluster (SkM). UMAP plots are colored by log-normalized transcript counts. **G** – UMAP plot showing expression of *Twist2* in the dermis fibroblast cluster. Plot is colored by log-normalized transcript counts. **H** – Feature plots showing expression of genes detected in the adipocytes precursor cluster (BApre). UMAP plots are colored by log-normalized transcript counts. **I** – Dotplot illustrating expression of selected genes in different somite derived cell type clusters (BApre=brown adipocyte precursors, BA=brown adipocytes, Dpre= dermis precursors, SkMpre=skeletal muscle precursors, SkM=skeletal muscle). Scale represents log-normalized transcript counts.

We next isolated dorsal tissues at the forelimb level at E13.5, E14.5, and E15.5 to perform scRNAseq to characterize early stages of BAT development (Figure 1B). For stages E13.5 and E14.5, the isolated dorsal tissues include epidermis, dermis, mesenchyme, interscapular brown fat and skeletal muscle. For the E15.5 time point, to enrich BAT precursors, we removed the epidermis and underlying dermis before dissociation. Using the inDrops workflow (Klein et al., 2015), we generated single-cell transcriptomes from at least 6000 cells per stage, resulting in a final dataset of 20,000 cells after quality control filtering (Supplementary figure 1A). Uniform Manifold Approximation and Projection (UMAP) based 2-dimensional embedding shows that cells from the different time points segregate based on transcriptional similarity rather than with embryonic age (Figure 1C, Supplementary figure 1B). Using Leiden clustering algorithm, we identified 22 clusters representing the somitic and non-somitic cell types that reside in the dorsal trunk region of the murine embryo (Figure 1D). Both Leiden and Louvain clustering algorithms generated similar cell clusters (Figure 1D, Supplementary figure 1C). We manually annotated the cell types corresponding to the different clusters using well-established marker genes for the various lineages. We merged cell clusters belonging to the same cell type to obtain a total of 18 cell clusters. Among somitic cell types, we identified cells that originate from the dermomyotome, including brown adipocytes (*Pparg*, *Cidea*, *Prdm16*, *Ucp1*), skeletal muscle (*Pax7*, *Myog*, *Ttn*, *Actn2*, *Myh3*) and dermal fibroblasts (*En1*, *Twist2*, *Dpt*, *Crabp1*) (Figure 1E-G, 1I). In addition, we could annotate smooth muscle cells (*Cnn1*, *Acta2*), muscle connective tissue (*Ngfr*, *Osr1*, *Osr2*) (Supplementary Figure 1E-F), cartilage (*Cnmd*, *Col2a1*), endothelial cells (*Cdh5*, *Kdr*) and meninges (*Foxc1*, *Cldn11*, *Aldh1a2*) (Supplementary Figure 1D, G-I). Interestingly, most of the somitic cells, except endothelial cells and skeletal muscle cells, lie closely together on the UMAP embedding suggesting that they have closely related transcriptional signatures (Figure 1D). Among non-somitic cells, we found macrophages (*Csf1r*, *C1qb*), mast cells (*Srgn*, *Kit*), neutrophils (*Ngp*, *Lccn2*), Schwann cells (*Sox10*, *Mpz*), and epidermal cells (*Krt5*, *Krt14*) (Supplementary Figure 1J-P). In conclusion, our single-cell transcriptomics analysis captured major somitic cell types including cells of the brown adipocyte lineage present in the mouse dorsal trunk during embryonic development.

### Identification of brown adipocyte precursors in the developing trunk

In addition to the differentiated cell types described above, our scRNAseq analysis also revealed precursor populations for several lineages. For instance, *Pax7*-expressing skeletal muscle precursors formed a distinct cell cluster which created a continuum to *Ttn*-expressing myocytes (Figure 1F, 1I). We also observed a cluster of cartilage precursors (Cpre) which expressed *Scx* and chondrogenic markers (*Sox9, Cnmd, Col2a1, Acan)* (Supplementary Figure 1D, 1I).

We also recognized a cluster containing prospective brown adipocyte precursors (BApre). Cells in this cluster express known brown adipocyte precursor markers such as *Pdgfra, Ebf2, Cd34* and *Pparg* ((Wang et al., 2014), Figure Figure 1E, H-I). This cluster includes cells from all 3 embryonic stages (Supplementary Figure 1B). In mouse, at E14.5, Ebf2 marks brown fat precursors but its expression is not restricted to these cells (Wang et al., 2014). In our scRNAseq dataset, we observed *Ebf2* mRNA expression in clusters of brown adipocytes, brown adipocyte precursors, connective tissue and precartilage but not in dermis (Figure 1H, Supplementary Figure 2A). Immunofluorescence analysis of E12.5 to E15.5 embryos confirmed the expression of Ebf2 in the presumptive brown fat, skeletal muscle and precartilage (Supplementary Figure 2A).

To better define the developmental trajectory of brown adipocyte precursors, we analyzed the single cell transcriptomes of different embryonic stages individually. UMAP-based embedding and Leiden clustering revealed similar somitic cell types which clustered together, except for skeletal muscle and endothelial cells, as in the combined temporal analysis (Supplementary figure 2B-D). Differential gene expression analysis identified *Gata6*, a transcription factor not previously associated with brown adipogenesis, as differentially expressed in the BAPre cluster (Figure 2A-D). Immunofluorescence analysis from E12.5 to E15.5 indicated that Gata6 starts to appear at E12.5 in the presumptive brown fat area under the dermis (Figure 2E). At E13.5, some of the Gata6 expressing cells also showed expression of Pparg at the mRNA (Figure 2D) and protein level (Figure 2F). Differentiating Pparg-positive immature adipocytes as well as Pparg-negative adipocyte precursors continued to express Gata6 at E14.5 (Figure 2G). At E15.5, Pparg-expressing adipocytes expressed *Gata6* mRNA (Figure 2C-D) but had downregulated the Gata6 protein except in the periphery of the BAT (Figure 2H). Downregulation of Gata6 in lipid-vesicle containing adipocytes at E15.5 suggests that Gata6 expression marks the brown adipocyte precursor state.

**Figure 2.**
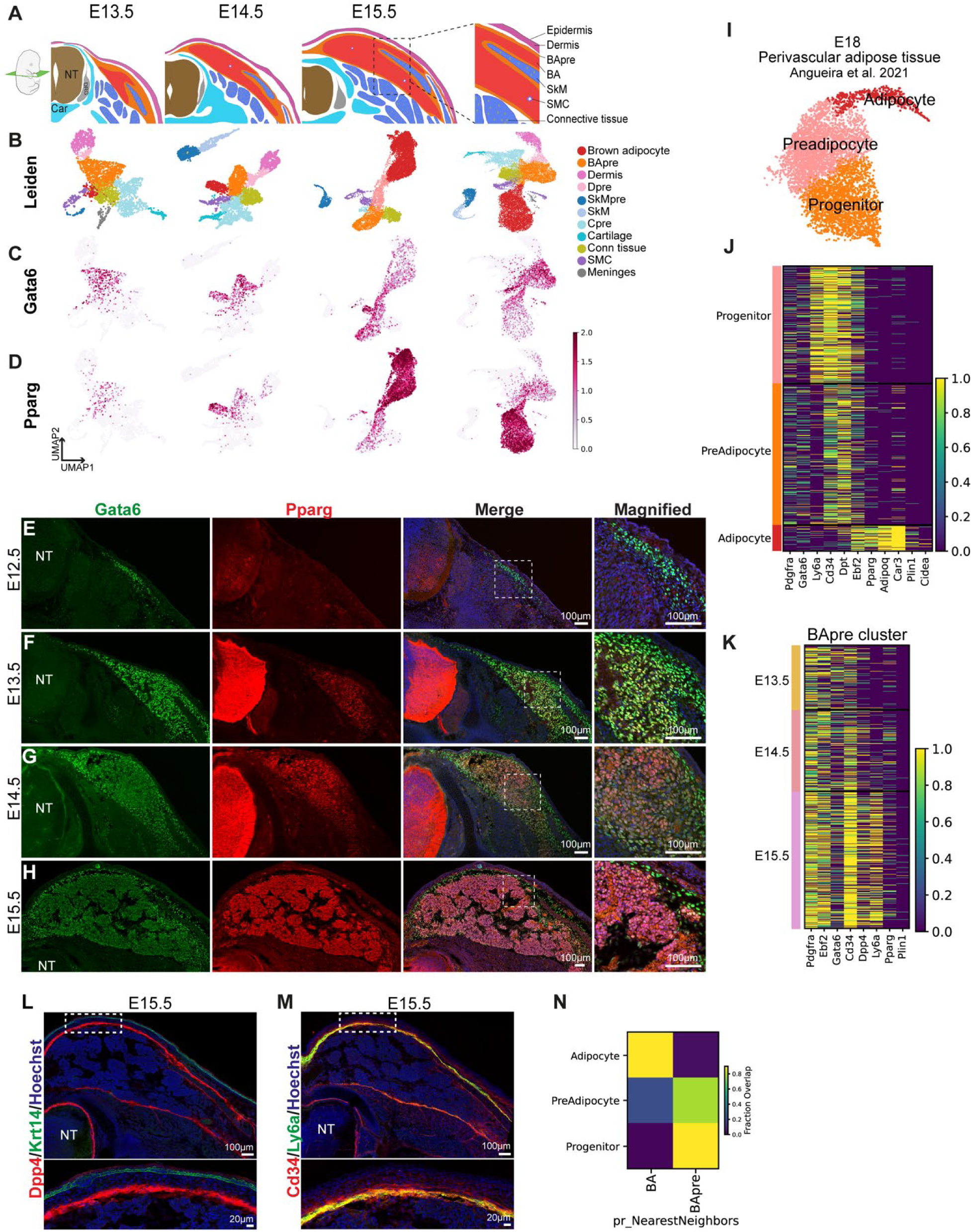
**A** – Schematic illustrating cell types identified and their respective location characterized by single cell RNA sequencing and immunofluorescence analysis in this study in the interscapular region of developing mouse embryo at E13.5-E15.5 stage. **B – D** – UMAP plots showing Leiden clusters representing somitic derivatives (B) and expression of *Gata6* (C) and *Pparg* (D) in the single cell transcriptomics data from developing mouse embryos. Scale represents log-normalized transcript counts. SMC=smooth muscle cells, SkM=skeletal muscle, BA= brown adipocytes. **E – H** – Immunofluorescence analysis of developing interscapular fat in mouse from embryonic day (E) 12.5 to 15.5. Images represents transverse section of mouse embryos at the forelimb level showing the interscapular region. Presence of Gata6 positive and Pparg negative cells detected by antibody staining at E12.5 (A). Expression domain of Gata6 expands and double positive cells are detected at E13.5 (B). Expression of Gata6 reduced at E14.5 and only detected in a few cells at E15.5 as majority of cells become Pparg positive (C, D). Note expression Gata6 in the hair follicle at E15.5 (D). NT= Neural tube. **I** – UMAP plot generated from mouse single cell transcriptomics data from the developing mouse perivascular adipose tissue on embryonic day 18 described in Angueira et al. 2021. Cells were clustered using Leiden clustering and cell clusters representing the adipocyte lineage were selected. See Supplementary Figure 2E for all clusters. **J** – Heatmap showing gene expression patterns in adipogenic cells sub-setted from the single cell transcriptomics data from the developing perivascular adipose tissue on embryonic day 18 described in Angueira et al. 2021. Scale represents log-normalized transcript counts. **k** – Heatmap showing expression of BApre specific markers at different time points. Scale represents log-normalized transcript counts. **L** – Transverse section at the forelimb level of a E15.5 embryo stained for Dpp4 and Keratin 14 (Krt14) antibody to detect cells from BApre cluster. NT= Neural tube. **M** –Transverse section from the interscapular region of a E15.5 embryo showing coexpression of Ly6a/Sca1 and Cd34 using antibody staining. NT= Neural tube. **N** – Classifier based machine-learning classification of perivascular adipose cells. A kNN-classifier trained on mouse single cell clusters was used to predict identities of cell clusters described in Angueira et al. 2021 study. Heatmaps depict fraction of mouse cluster assignments for adipocyte, preadipocytes and progenitors identified in the perivascular adipose tissue. BA= brown adipocytes and BApre=brown adipocyte precursors identified in this study.

Angueira et al. (2021) recently identified progenitors that give rise to brown adipocytes in mouse periaortic (pVAT) brown adipose cells analyzed by scRNAseq at E18 and perinatal stages (Angueira et al., 2021). We reanalyzed their E18 dataset which contains 16,967 single cell transcriptomes. Following UMAP embedding and Leiden clustering, we identified 21 clusters representing the cell types described in the original study (Supplementary Figure 2E). After manual annotation of the adipocyte clusters, we isolated three clusters which correspond to adipocytes (*Cidea*), preadipocytes (*Pparg*) and adipocyte progenitors (*Cd34*) (Figure 2I, Supplementary Figure 2F). Remarkably, we observed expression of *Gata6* in the adipocyte progenitors cluster (Figure2J, Supplementary Figure 2F). *Pi16*, *Cd34* and *Ly6a*, which were also identified as brown adipocyte progenitor markers in this report (Figure 2J) were also differentially expressed in the BApre population identified in our study (Figure 2K). Immunofluorescence analysis at E15.5 confirmed the expression of the adipocyte precursors markers Gata6, Dpp4, Cd34 and Ly6a at the periphery of the developing brown adipose tissue (Figure 2H, L-M). Their location suggested that they might correspond to precursor cells involved in the generation of mature brown fat. To further analyze the overlap between the brown adipocyte progenitor cell population identified in the pVAT dataset and our BApre cluster, we trained a k-NN classifier on the brown adipocyte (BA) and BApre cluster identified in our dataset. This analysis revealed a high similarity between the pVAT progenitor cluster and our BApre cluster suggesting that pVAT undergoes a similar developmental sequence to that of the interscapular brown fat (Figure 2N).

### Gata6 marks early brown adipose tissue precursors

Next, to better characterize the differentiation trajectory of cells of the BApre cluster, we performed RNA velocity analysis on E14.5 single-cell transcriptomics data, restricting our analysis to somitic cell types (Figure 3A, Supplementary Figure 3A). This identified a differentiation trajectory from the BApre cluster to the brown adipocyte cluster (Figure 3A-B). Phase portraits for genes such as BApre specific *Gata6*, or BA specific *Car3* showed positive velocity in BApre and BA cluster respectively. This further supports the notion that *Gata6*-expressing cells of the BApre cluster differentiate into brown adipocytes.

**Figure 3.**
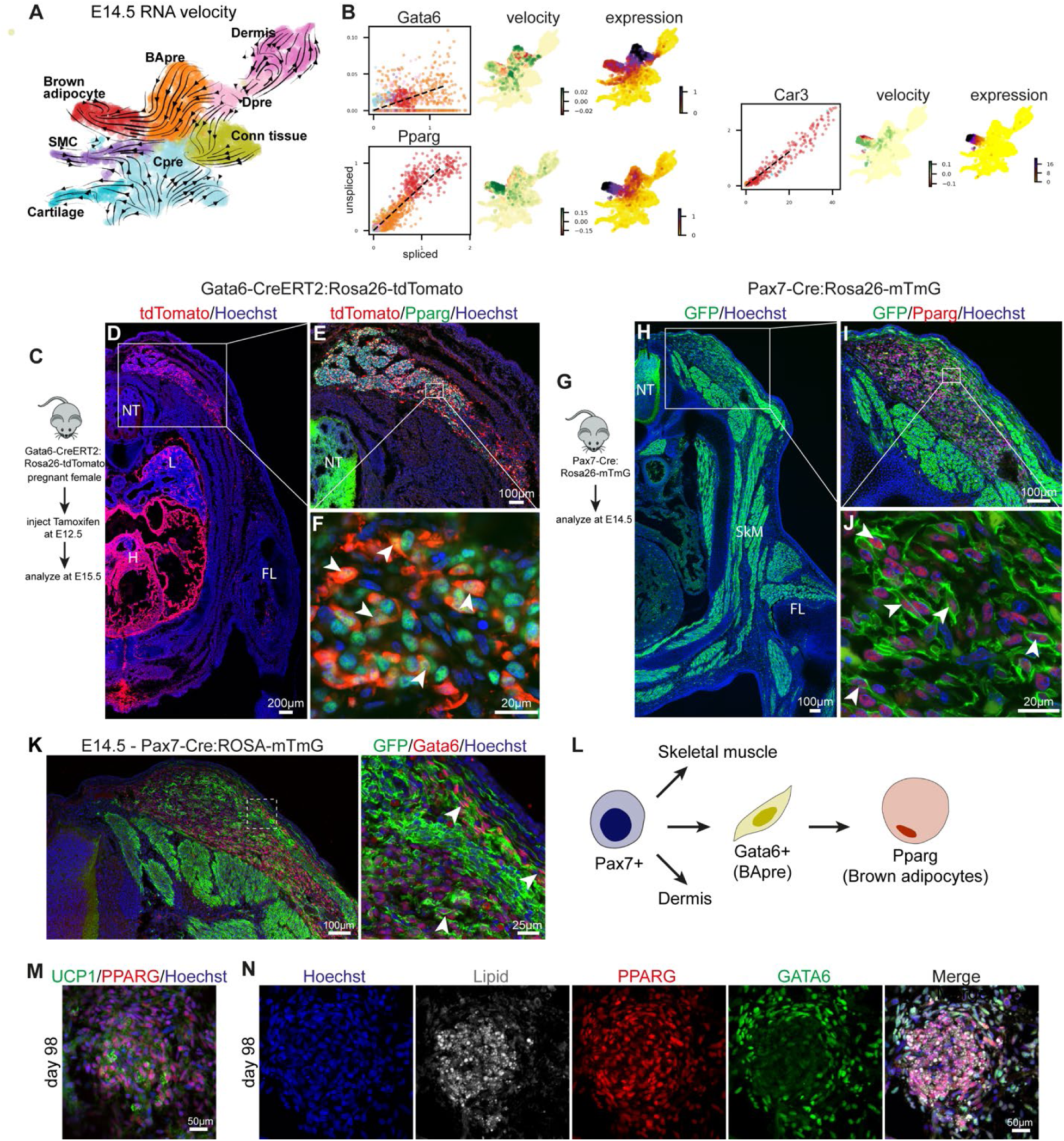
**A** – RNA velocity analysis of embryonic day 14.5 single cell transcriptomes from mouse embryo. For simplicity cell clusters representing somitic lineages were subclustered. Velocity field are projected onto the UMAP plot and arrows show the local average velocity. Cell types are clustered using Leiden clustering. BApre=Brown adipocyte precursor, SMC= smooth muscle cell, Dpre=dermis precursors, Conn tissue= Connective tissue cells, Cpre=Cartilage precursor and Cartilage). **B** – Velocity and Expression plots of *Gata6*, *Pparg* and *Car3* on E14.5. **C** – Schematic illustrating the lineage tracing experiments in Gata6-CreERT2:Rosa26-tdTomato mice. A single dose of Tamoxifen was injected into pregnant females at E12.5 and embryos were harvested at E15.5. **D** – Transverse section at the forelimb level of E15.5 embryos were stained for anti-RFP antibody to detect tdTomato positive cells. Gata6 progeny was detected in the interscapular brown adipose tissue (BAT), the lung (L) and the heart (H). NT= Neural tube. **E** – Magnified image of the transverse section from the interscapular region showing brown adipose tissue stained with Pparg and RFP (tdTomato) antibody. **F** – Higher magnification image of brown adipocytes in the interscapular region showing double positive cells for Pparg and tdTomato. Arrowheads mark some of the double positive cells. **G** – Schematic illustrating lineage tracing strategy to label Pax7 progeny during mouse development. Pax7-Cre:Rosa26-mTmG embryos were analyzed at E14.5. **H** – Representative image showing Pax7 progeny at E14.5. Transverse section at the forelimb level of E14.5 embryos were stained for anti-GFP antibody to detect membrane GFP positive cells. GFP labelling identifies Pax7 progenies in the interscapular brown adipose tissue and the skeletal muscle bundles (SkM) in the dorsal and limb region. NT= Neural tube. **I** – Transverse section of E14.5 Pax7-Cre:Rosa26-mTmG embryo showing interscapular region stained with antibodies against Pparg and GFP to label brown adipocytes and Pax7 progeny. **J** – Representative high magnification image showing double positive cells stained for nuclear Pparg and membranous GFP. Arrowheads mark double positive cells. **K** – Transverse section of E14.5 Pax7-Cre:Rosa26-mTmG embryo showing interscapular region stained with antibody against GFP to label Pax7 progeny and Gata6 to label brown adipocyte precursors. Inset shows magnified view. Arrowheads mark some of the double positive cells. NT= Neural tube. **L** – Schematic illustrating proposed model of interscapular brown fat differentiation in mouse. Pax7 positive (Pax7+) dermomyotomal cells give rise to Gata6 expressing (Gata6+) brown adipocyte precursors (BApre) that differentiate into brown adipocytes. **M** – Immunofluorescence analysis of human fetal brown adipose tissue isolated from the scapular region of a 98-day old fetus. Immunofluorescence analysis confirming expression of UCP1 and PPARG in the tissue. **N** – Immunofluorescence analysis of consecutive section of human fetal brown adipose tissue from a 98-day old fetus shown in Figure 3L. Antibody staining showing expression of GATA6 in the brown adipocytes precursors and PPARG expression in the precursors and lipid droplets containing adipocytes.

To experimentally demonstrate that Gata6 is expressed by precursors which give rise to brown adipocytes, we used a Gata6-CreERT2:Rosa26-tdTomato mouse line to label Gata6-expressing cells during embryonic development (Donati et al., 2017). We induced recombination using tamoxifen injection at E12.5 and analyzed the contribution of labelled cells to brown adipose tissue at E15.5 (Figure 3C). Immunofluorescence analysis for tdTomato, which labels the descendants of Gata6-expressing cells confirmed that at E12.5, they give rise to heart and lung bud cells as expected (Figure 3D). Among the somite-derived cells, tdTomato positive cells were only found in the interscapular BAT region and not in other somite derivatives such as skeletal muscle, cartilage or endothelial cells. TdTomato-positive cells in the interscapular BAT also stained positive for Pparg at E15.5 confirming that these cells are adipocytes (Figure 3E-F). We also found rare tdTomato cells closely associated to Myh11 expressing smooth muscle cells next to the blood vessels in the interscapular brown adipose tissue (Supplementary Figure 3B). These cells could correspond to the recently identified brown adipocyte precursors deriving from the vascular smooth muscle lineage (Shamsi et al., 2021). In conclusion, the lineage tracing analysis confirmed that Gata6-expressing cells present at E12.5 in the mouse dorsal trunk mostly represent a brown adipocyte precursor population.

To further investigate the origin of these Gata6 precursors, we used a Pax7-Cre:Rosa26-mTmG mouse line to analyze the contribution of Pax7 descendants to Gata6-positive brown fat precursors tissue at E14.5 (Figure 3G) (Hutcheson et al., 2009). As expected, dorsal, ventral and limb skeletal muscle as well as Pparg-expressing brown adipocytes were identified as Pax7 descendants (Figure 3H-J). Membrane GFP also labeled Gata6 positive cells in the brown fat region indicating that these adipocyte precursors derive from an earlier Pax7-expressing population (Figure 3K). Taken together, our experiments indicate that Pax7-positive somitic progenitors give rise to a Gata6-positive precursor population that differentiates into brown adipocytes (Figure 3L).

To further examine whether Gata6 expression is conserved in human brown adipogenesis, we performed immunofluorescence analysis on 98 and 135 day old (estimated post-conceptual age) human fetal tissue from the interscapular and scapular regions. We detected expression of EBF2 and GATA6 in cells adjacent to skeletal muscle tissue (detected by MYHC antibody) under the dorsal dermis and dispersed between the deeper dorsal muscle bundles at 98 days of development (Supplementary figure 3C). As in mouse, EBF2 was expressed in a broader territory in the dermis and deeper muscle areas. We also observed rare UCP1 expressing lipid-filled PPARG positive adipocytes in the deeper muscle area (Figure 3M, Supplementary Figure 3D). In these regions, we detected PPARG and GATA6 double-positive cells, supporting the existence of GATA6-positive brown adipocyte progenitors in humans (Figure 3N, Supplementary Figure 3E). Brown adipocytes identified by the presence of lipid droplets, expression of PLIN1 and high expression of PPARG did not express GATA6 suggesting that GATA6 also marks an early stage of brown adipocyte development in humans.

### Recapitulation of human brown fat development in vitro from pluripotent stem cells

We previously reported that iPSCs can be induced to form paraxial mesoderm and subsequently differentiated into skeletal muscle in vitro (Chal et al., 2016; Chal et al., 2015; Diaz-Cuadros et al., 2020). As brown adipocytes originate mostly from paraxial mesoderm and share a common precursor with skeletal muscles in the dermomyotome, we first aimed to recapitulate the development of Pax-3 positive dermomyotomal cells in vitro. To achieve this, we treated human iPSCs with the Wnt agonist CHIR and the BMP inhibitor LDN in serum-free conditions to differentiate them to presomitic mesoderm (PSM) (Figure 4A). At day 2-3, most of the cells expressed the PSM markers MSGN1 (95±0.4%) and TBX6 (Figure 4B-C, Supplementary Figure 4A) indicating that they acquired a PSM fate. To further differentiate the PSM cells into dermomyotomal cells, we treated cultures with IGF, FGF, HGF, and LDN (Chal et al., 2016; Chal et al., 2015). To monitor cell differentiation towards the dermomyotome fate, we used CRISPR-Cas9 to knock-in a Venus fluorescent reporter into the PAX3 locus in the NCRM1 iPSC line (Supplementary figure 4B). We used this reporter line to track the appearance of PAX3 in differentiating cells in vitro. On day 8, differentiating cells started to express PAX3, indicating the appearance of somitic/dermomyotomal cells (Figure 4D). 45±6.7 % of the cells in the culture were PAX3-positive (Figure 4E, Supplementary Figure 4C). We further differentiated these cultures in medium containing HGF and IGF as described previously (Chal et al., 2016; Chal et al., 2015). Using a PAX7-Venus reporter iPSC line, we observed that at day 14-16, cultures harbored large populations of non-overlapping PAX7-expressing and EBF2-expressing cells (Figure 4F) (Al Tanoury et al., 2020).

**Figure 4.**
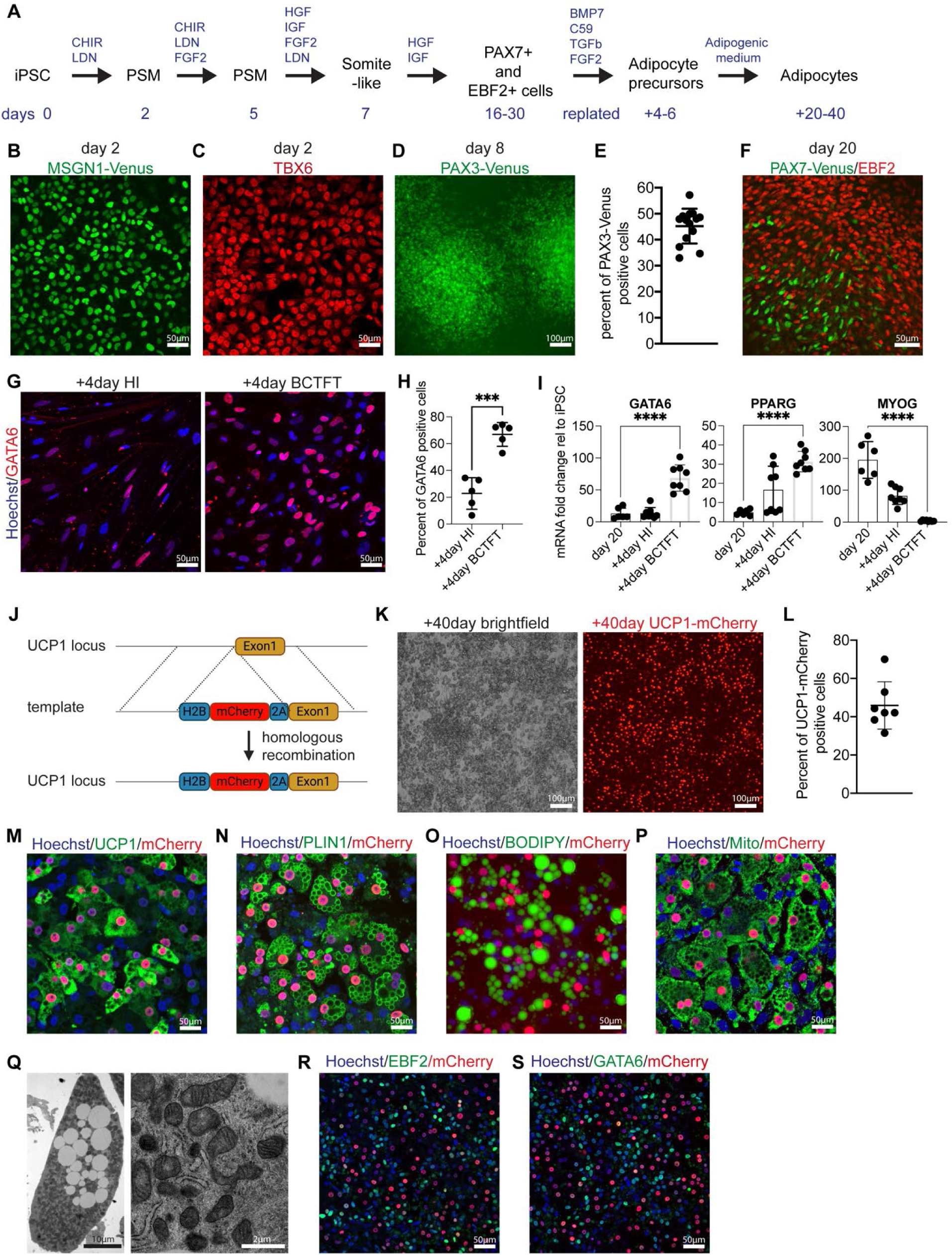
**A** – Schematic describing the steps to differentiate human induced pluripotent stem cells (iPSCs) into presomitic mesoderm (PSM), followed by a somite-like stage and somitic derivatives in a monolayer culture. Cells are replated between day 16-30 of differentiation to generate brown adipocytes. + indicates additional days after replating. **B** – Venus fluorescence signal in MSGN-Venus knock-in iPSC line on day 2 of differentiation. **C** – Immunofluorescence analysis for transcription factor TBX6 on day 2 of differentiation. **D** – Venus fluorescence signal in PAX3-Venus knock-in iPSC line on day 8 of differentiation. **E** – FACS analysis quantification of PAX3-venus positive cells on day 8 of differentiation. Mean ± SD, n=14. **F** – Immunofluorescence analysis of transcription factor EBF2 and PAX7 on day 20 of differentiation. **G** – Immunofluorescence analysis for transcription factor GATA6 after replating and culturing of 20-day old cultures in HI medium or adipogenic BCTFT medium for 4 days (+4 represents 4 days after replating). **H** – Quantification of GATA6 positive cells in figure panel G. Mean ± SD, n=5, t-test, p≤0.001. **I** – RT-qPCR analysis for *GATA6*, *PPARG* and *MYOG* on day 20 of differentiation and 4 days after replating in HI medium or adipogenic BCTFT medium (+4 represents 4 days after replating). Mean ± SD, n=6-8, t-test, p≤0.0001. **J** – Diagram outlining the targeting strategy used to generate UCP1-mCherry knock-in iPSC line. **K** – Brightfield and mCherry signal in iPSC-derived brown adipocyte cultures on day 40 after replating. **L** – Quantification of mCherry positive cells in iPSC-derived brown adipocyte cultures on day 40 after replating. Mean ± SD, n=7. **M**–**N** – Immunofluorescence staining for UCP1 (J), PLIN1 (K) in the UCP1-mCherry knock-in iPSC line derived brown adipocyte cultures on day 40 after replating. **O** – Neutral lipid staining using 0.5mM BODIPY in iPSC-derived brown adipocyte cultures on day 40 after replating. **P** – Immunofluorescence staining for mitochondria on iPSC-derived brown adipocyte cultures on day 40 after replating. **Q** – Representative transmission electron micrographs demonstrating ultrastructural characteristics (left) and mitochondrial characteristics (right) of iPSC-derived brown adipocyte. **R**–**S** – Immunofluorescence staining for EBF2 (O), GATA6 (P) in the iPSC-UCP1mCherry knock-in iPSC-derived brown adipocyte cultures on day 40 after replating.

To divert the differentiation of the paraxial mesoderm cultures towards an adipogenic fate, we next sought to identify potential signaling pathways involved in brown adipogenesis. We performed differential gene expression analysis with the BApre cluster of the mouse dorsal trunk scRNAseq dataset (Supplementary Table1). Using KEGG pathway analysis, we identified several pathway-specific genes differentially expressed in the BApre cluster (Supplementary Figure 4D). This included genes involved in BMP signaling (*Bmp7*, *Bmp4*, *Bmpr1a*, *Bmpr2*), TGFbeta signaling (*Tgfbr2*, *Gdf10*), Thyroid hormone signaling (*Creb3l1*,*Gnas*, *Gpx7*, *Creb5*, *Hsp90b1*), Wnt signaling antagonists (*Sfrp1*, *Sfrp2*, *Sfrp4*, *Dkk2*) and FGF/PI3-AKT pathway (*Akt1*, *Igf1*, *Fgfr1*) (Supplementary Figure 4E-G). Thus, to induce the brown adipocyte precursor fate, we dissociated the primary cultures between day 16-30 of differentiation and replated the cells in a medium containing BMP7, the Wnt antagonist C-59, TGFb1, FGF2 and Triiodothyronine (BCTFT medium) for 4-6 days (Figure 4A). Strikingly, after culturing cells in the BCTFT medium we saw a strong increase in number of GATA6-expressing cells compared to cells cultured in medium containing HGF and IGF (Figure 4G-H). The cells also showed an upregulation of *GATA6* and *PPARG* and a downregulation of *MYOG1* mRNA, suggesting that the BCTFT medium can induce an adipogenic fate while repressing the myogenic fate (Figure 4I).

To further drive the differentiation of GATA6-positive adipocyte precursors into lipid vesicles containing adipocytes, after 4-6 days in BCTFT medium, replated cells were switched to a new medium containing an adipogenic cocktail including 3-Isobutyl-1-methylxanthine, Ascorbic acid, Triiodothyronine, TGFb inhibitor SB431542, Dexamethasone, EGF, Hydrocortisone and Rosiglitazone for another 20-40 days (Figure 4A) (Hafner et al., 2016). To better follow the differentiation of iPSCs into brown adipocytes, we generated a UCP1-mCherry fluorescent human iPSC reporter line using CRISPR-Cas9. In this knock-in line, mCherry is cleaved from UCP1 during translation and targeted to the nucleus using a H2B tag (Figure 4J). In replated cultures examined after 20 days, lipid-filled adipocytes started to appear and the adipocyte number steadily increased over the following 20 days. By day 40 of differentiation, the replated cultures contained 46±12% UCP1-mCherry positive cells (Figure 4K-L). Immunofluorescence staining demonstrated colocalization of UCP1 protein in adipocytes with UCP1-mCherry expression (Figure 4M). PLIN1 and BODIPY staining confirmed that iPSC-derived brown adipocytes (iPSC-BAs) contain multilocular lipid droplets (Figure 4N-O). We further confirmed expression of bona fide brown adipocyte markers such as *UCP1, PPARG, CIDEA* and *PLIN1* using RT-qPCR in replated cultures after 40 days (Supplementary figure 4H).

iPSC-BAs contain abundant mitochondria (a characteristic of brown adipocytes) as revealed by immunofluorescence for mitochondrially encoded cytochrome c oxidase II (MT-CO2) and electron microscopy analysis at 40 days after replating (Figure 4P-Q). These cultures contained a fraction of UCP1-negative cells, which expressed EBF2 and/or GATA6 cells suggesting these cells might correspond to brown adipocyte precursor cells (Figure 4R-S). Dissociation of the replated cultures at 40 days resulted in loss of the UCP1 expressing lipid filled adipocytes. However, when replated, these dissociated cultures were able to produce large amounts of UCP1-positive cells indicating that a significant proportion of the UCP1-negative cells are brown adipocyte precursors. Periodic acid staining and electron microscopy showed that at this stage, many cells accumulate glycogen, a characteristic of differentiating brown adipocytes (Supplementary Figure 4I-J) (Mayeuf-Louchart et al., 2019). Thus, our protocol recapitulates a developmental trajectory similar to that observed for brown fat differentiation in mouse embryos, leading to efficient differentiation of human iPSCs into UCP1-expressing brown adipocytes under serum free culture conditions.

### scRNAseq analysis of the human brown adipogenic lineage differentiated in vitro

To further characterize the differentiation of iPSCs into adipocytes, we performed scRNAseq of the cultures at day 6, 10, 15, 20 and 30 before replating and 20 days and 40 days after replating (Figure 5A). For each time point we analyzed at least 2000 cells. After quality control and filtering, we analyzed 14,313 cells from the cultures before replating and 6,676 cells from the replated cultures (Figure 5B-C). Using UMAP embedding and Leiden clustering algorithms, we identified 12 clusters in the non-replated cultures which were annotated based on well-established cell identity markers (Figure 5D). At day 6, we only identified one cluster corresponding to the anterior presomitic mesoderm stage characterized by expression of marker genes such as *TBX6* and *TCF15* (Figure 5D, Supplementary Figure 5A). By day 10 the cultures expressed the dermomyotomal markers *PAX3* and *TCF15* as well as sclerotomal ones such as *PAX9*, *NKX3.2* and *FOXC2* indicating that cells have reached the differentiating somite stage (Figure 5D, Supplementary Figure 5A-B). At day 20 and 30, several distinct somite-derived cell types such as *PDGFRA* expressing fibroblasts, *EBF2*/*GATA6* expressing brown adipocyte precursors, *CNN1* positive smooth muscle cells, *PAX7* expressing muscle precursors, *TTN* expressing skeletal muscle and *SOX9* expressing cartilage precursors were identified in the cultures (Figure 5D, 5F, 5H, Supplementary Figure 5A). In addition, we identified a smaller contingent of differentiating neural cells which are lineages also deriving from the neuro-mesodermal precursors first generated in the differentiation protocol (Figure 5D, Supplementary Figure 5A-B) (Diaz-Cuadros et al., 2020). Analysis of the replated cultures at 20 and 40 days identified *PDGFRA*-expressing fibroblasts, *EBF2*/*GATA6*-expressing adipocyte precursors and *PPARG*-expressing adipocytes as well as small populations of skeletal muscle, cartilage and neural cells (Figure 5G, 5I, Supplementary Figure 5C-D). As single cell dissociation disrupts lipid filled adipocytes, the number of *UCP1*-positive adipocytes was small, but we were nevertheless able to capture some cells expressing *CIDEA* and *UCP1*, which represent mature brown adipocytes (Figure 5I, Supplementary Figure 5C).

**Figure 5.**
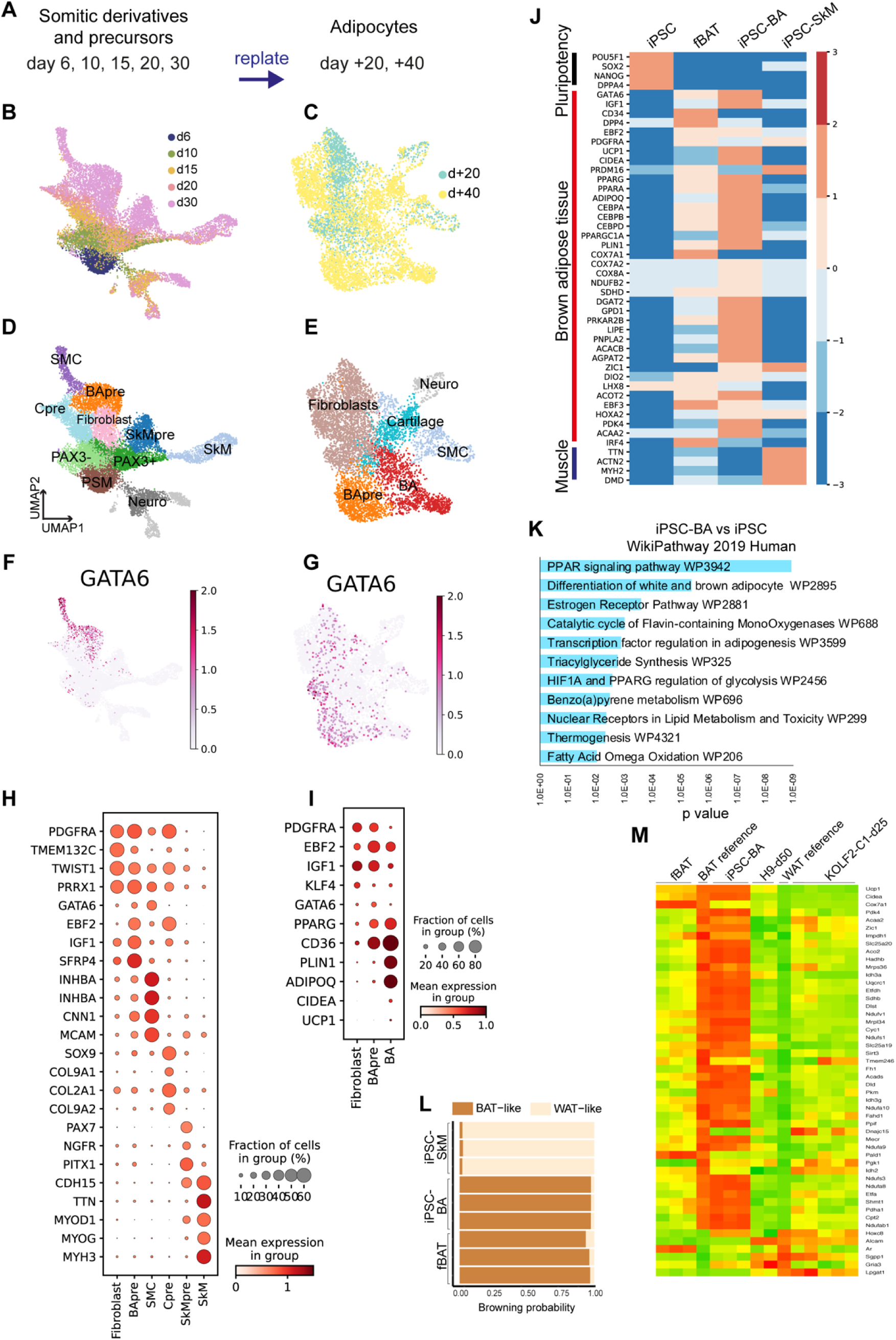
**A** – Scheme showing collection of differentiating iPSC samples for single cell RNA sequencing analysis. Cells were differentiated into somitic derivatives for 30 days and cells were collected on day 6, 10, 15, 20 and 30 days of differentiation. At day 20-30 cells were dissociated and replated for secondary differentiation in adipogenic conditions and cells were harvested on day 20 and 40 after replating (+ represents days after replating). **B** – **C** – UMAP embedding of the single cells isolated from iPSC cultures after day 6 - day 30 (14,313 cells) (B) and day 20 - day 40 (6,676 cells) after replating (C). Bkknn batch correction was performed with 50 PC dimensions. Colors indicate days in differentiation. **D** – **E** – UMAP plot showing cell clusters identified in differentiating iPSCs using Leiden clustering on day 6-day 30 cultures (D) and day 20-day 40 replated cultures (E). PSM=anterior presomitic mesoderm, PAX3+=PAX3 expressing dermomyotomal cells, PAX3-=sclerotomal cells, SkMpre=Skeletal muscle precursor, SkM=Skeletal muscle, SMC=Smooth muscle cells, BA=Brown adipocytes, BApre= brown adipocyte precursors, Neuro=neural cells. **F** – **G** – UMAP plots showing expression pattern of brown adipocyte precursor marker *GATA6*. Scale represents log-normalized transcript counts. **H** – Dot plot showing differentially expressed genes identified in differentiated cell clusters Fibroblast, BApre, SkM, SkMpre, Cpre, SMC. Scale bar represents log-normalized transcript counts. **I** – Dot plot showing differentially expressed genes identified in differentiating adipogenic cultures. BApre=brown adipocyte precursors, BA=brown adipocyte. Scale bar represents log-normalized transcript counts. **J** – Heatmap showing differentially expressed genes in undifferentiated human pluripotent stem cell (iPSC), human fetal brown adipose tissue (fBAT), 40 day old replated iPSC-derived adipogenic cultures (iPSC-BA) and iPSC derived skeletal muscle (iPSC-SkM, described in Al Tanoury et al. 2021). Scale bar represents log normalized read counts. **K** – WikiPathway 2019 Human analysis of top 200 differentially expressed genes (FDR<0.05, log2|FC|>6) in iPSC-BAs compared to undifferentiated iPSCs. **L** – Estimation of browning probability of iPSC derived brown adipocytes (iPSC-BA), iPSC derived skeletal muscle (iPSC-SkM) and human fetal brown adipose tissue (fBAT) using PROFAT database. Numbers represent replicates. **M** – Heatmap comparing transcriptomic profiles of iPSC derived brown adipocytes (iPSC-BA), human fetal brown adipose tissue (fBAT), human embryonic stem cell derived brown adipocytes in Zhang et al. 2020 (H9-d50) and iPSC derived brown adipocytes in Carobbio et al. 2021 (KOLF2-C1-d25).

To molecularly characterize the iPSC-BAs which are mostly lost during dissociation in the scRNAseq analysis, we performed bulk mRNA sequencing of undifferentiated iPSCs and replated cultures at 40 days. In parallel, as a reference for differentiated adipocytes, we analyzed human fetal BAT (fBAT) isolated from 115, 122 and 125-day post-conceptual age fetuses from the interscapular and scapular regions. As an outgroup, we used bulk RNAseq data from iPSC-derived skeletal muscle cultures (iPSC-SkM) (Al Tanoury et al., 2021). Brown adipocyte-specific genes such as *UCP1*, *CIDEA*, *PARGC1A* and *DIO2* were specifically expressed in iPSC-BAs and human fetal BAT (Figure 5J). Markers for precursors identified in this study such as *GATA6* and *DPP4* were also detected suggesting the presence of adipocyte precursors in differentiating cultures (Figure 5J). The top 200 differentially expressed genes in iPSC-BAs corresponded to human WikiPathways related to ‘PPARG signaling pathway’, ‘Estrogen receptor pathway’ and ‘Thermogenesis’ (Figure 5K). When analyzed using the ProFat database (Cheng et al., 2018), the browning probability of iPSC-BAs was comparable to fetal brown adipocytes (Figure 5L). Recently, two studies have described the directed differentiation of brown adipocytes from hPSCs (Carobbio et al., 2021; Zhang et al., 2020). Comparison of our iPSC-BAs with the terminally differentiated adipocytes from the two studies – 50 day old adipocytes from Zhang et al. (H9-d50) and 25 day old adipocytes from Carobbio et al. (KOLF2-C1-d25) showed that cells differentiated according to our protocol exhibit a higher degree of similarity to the human fetal BAT and brown adipocytes gene expression profile (Figure 5M, Supplementary Figure 5E).

### iPSC derived brown adipocytes respond to beta-adrenergic stress

Next, we sought to investigate the functional potential of iPSC-BAs generated in vitro. Generally, brown adipocytes are characterized by their potential to generate heat in response to beta-adrenergic stimulus and downstream activation of the cAMP pathway (Wang and Seale, 2016). To test the capacity of iPSC-BAs to respond to adrenergic stress, we used forskolin in 40 day-old replated adipogenic cultures. Cells treated with forskolin for 4 hours show increased *UCP1* mRNA levels compared to vehicle-treated cells (Figure 6A). Adrenergic stimulus also induces lipolysis of stored triglycerides into fatty acids and glycerol. To evaluate the lipolytic activity of the cells, we measured the amount of glycerol released in the medium after forskolin treatment. In comparison to vehicle control treated cells, forskolin-treated cells showed an increased glycerol release indicating that cells underwent lipolysis (Figure 6B). Brown adipocytes can utilize free fatty acids to induce proton leak and perform thermogenesis. To measure the ability of iPSC-BAs to generate heat, we used ERThermAC, a thermosensitive vital fluorescent dye (Wang et al., 2020). As temperature increases, the cells display lower ERthermAC fluorescence intensity. We preincubated iPSC-BAs with ERThermAC, treated with PBS or forskolin and measured the fluorescence intensity of the dye over time. We observed a reduction in dye intensity in forskolin-treated cells compared to vehicle-treated cells (Figure 6C). In contrast, no change in intensity was observed for BA precursors in either condition (Figure 6C). Overall, our data demonstrate that iPSC-BAs induce UCP1 expression, undergo lipolysis and generate heat in response to beta adrenergic stress.

**Figure 6.**
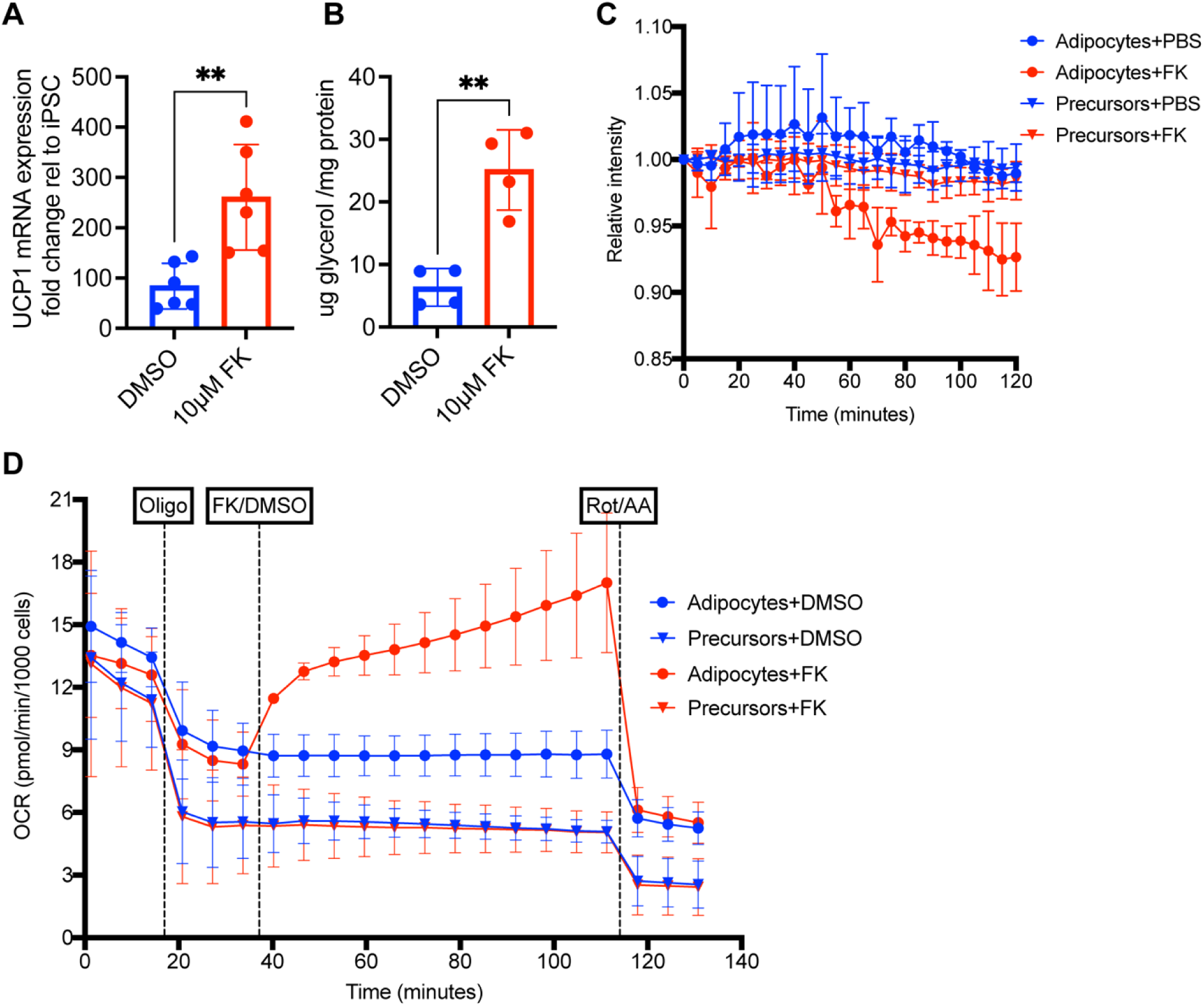
**A** – RT-qPCR analysis for *UCP1* on 40 day old replated iPSC-derived adipogenic cultures treated with 10µM forskolin or vehicle control DMSO treatment for 4 hours. Relative gene expression is shown as relative change to undifferentiated iPSC. Mean± SD, n= 5, t-test, p≤0.01. **B** – Measurement of glycerol released in culture medium 40 day old replated iPSC-derived adipogenic cultures treated with 10µM forskolin or vehicle control DMSO treatment for 4 hours. Glycerol amount normalized using total protein amount. Mean± SD, n= 4, t-test, p≤0.01. **C** – Measurement of fluorescent intensity of thermosensitive ERthermAC dye in iPSC derived precursors on day 20 of differentiation and brown adipocytes after 40 days of replating in response to 10µM forskolin or vehicle control DMSO. Each data point represents mean ± SD, n= 3. **D** – Seahorse analysis to measure oxygen consumption rate in iPSC derived precursors on day 20 of differentiation and brown adipocytes after 40 days of replating in response to 1.5µM oligomycin, 10µM forskolin or vehicle control DMSO and 1µM Rotenone and Antimycin (Rot/AA), respectively. Adipocytes or precursor cells were in XFe96 cell culture plates one week prior to the assay. Assay medium was prepared using Seahorse XF DMEM supplemented with 1 mM pyruvate, 2 mM glutamine and 10 mM glucose. Oxygen consumption rate was normalized by number of cells per well. Each data point represents mean ± SD, n= 4.

The process of thermogenesis is driven by the flow of protons through UCP1 in the mitochondrial inner membrane. To examine whether forskolin induced thermogenesis in iPSC-BAs is coupled to proton leak in mitochondria, we performed metabolic analysis of 40 day-old replated cultures. We measured the oxygen consumption rate (OCR) of iPSC-BAs with a Seahorse metabolic flux analyzer (Figure 6D). Upon treatment with Oligomycin, an ATP synthase inhibitor, the OCR in cells decreased. Interestingly, when iPSC-BAs were treated with forskolin together with Oligomycin, the OCR increased suggesting that proton leak is increased, leading to higher oxygen consumption (Figure 6D). In contrast, the precursors showed lower basal respiration level and did not show increased OCR upon forskolin treatment. As expected, inhibiting the electron transport chain using Rotenone (mitochondrial complex I inhibitor) or Antimycin (mitochondrial complex III inhibitor), resulted in a drop of OCR (Figure 6D). These analyses demonstrate the functionality of human iPSC-BAs differentiated in vitro.

## DISCUSSION

Studying brown fat lineage development has proven complicated due to the difficulty of identifying intermediate stages in differentiation. Single cell sequencing techniques together with lineage tracing methods now provide powerful approaches for exploring cell states during differentiation programs. In this report, we first characterized the development of interscapular mouse brown fat using scRNAseq. We analyzed the development of paraxial mesoderm-derived cell populations in the dorsal trunk at the forelimb level of mouse embryos which contains the interscapular brown fat. This analysis identified a previously unrecognized cell population corresponding to brown adipocyte precursors, characterized by expression of the transcription factor Gata6. Using RNA velocity and lineage tracing in mouse, we show that these cells derive from Pax7-positive precursors and give rise to Pparg-positive preadipocytes which differentiate into UCP1-positive brown adipocytes (Figure 3L).

This scRNAseq analysis also allowed the identification of specific signaling pathways associated to the Gata6-positive brown adipocytes’ progenitor state. We show that Bmp4 and Bmp7 and Bmpr1a and 2 are enriched in the Gata6-positive precursors. This is consistent with the well-established role of Bmps in the promotion of brown adipogenesis (Schulz et al., 2013; Tseng et al., 2008). We also observed transient expression of the Wnt inhibitors Sfrp1,2 and 4 and Dkk2 in the Gata6-positive precursors, consistent with the well-known anti-adipogenic role of Wnt signaling (Bagchi and MacDougald, 2021; Longo et al., 2004). The thyroid hormone and the FGF/IGF1/PI3K pathways which play a role in the control of thermogenesis and brown adipose development and physiology were also activated in the Gata6 precursors (Mullur et al., 2014; Ohta and Itoh, 2014). We used these signaling cues to modify a protocol previously established to differentiate human paraxial mesoderm in vitro (Chal et al., 2016). This allowed highly efficient generation of functional human brown adipocytes in vitro using defined and serum-free culture conditions. ScRNAseq analysis of the human cells differentiating in vitro show that they recapitulate a developmental sequence very similar to that observed in mouse in vivo. This sequence results in the production of GATA6-positive brown adipocyte precursors and then to PPARG-positive preadipocytes and finally to UCP1-positive adipocytes which exhibit functional properties characteristic of brown adipocytes.

Our scRNAseq analysis shows that brown adipocyte precursors tightly cluster with other somite-derived mesenchymal lineages such as dermis, muscle connective tissue, cartilage precursors and smooth muscles suggesting that they are transcriptionally closely related with these fibroblastic populations. In contrast, the skeletal muscle cluster is clearly segregated from the mesenchymal cluster in our dataset. This is consistent with the early divergence of the Pax7-expressing multipotent progenitors into skeletal muscle lineage by E11.5 revealed by lineage tracing experiments in mouse (Lepper and Fan, 2010). Lineage tracing analyses in mouse demonstrated that interscapular brown fat derives from Pax3/Myf5+ and Pax3/Pax7/Myf5+ cells (Sebo et al., 2018). Our lineage tracing experiments confirm that some of the mouse interscapular brown fat derives from Pax7-expressing precursors (Lepper and Fan, 2010; Sanchez-Gurmaches and Guertin, 2014) which can give rise to a Gata6-positive brown adipocyte precursor population. These early Pax7 precursors could correspond to the En1-positive cells of the central dermomyotome previously identified in mouse (Atit et al., 2006). The relatively low contribution of Gata6-tdTomato labeled cells to the adipose tissue could result from waves of adipocytes precursors differentiation as suggested by En1 lineage tracing (Atit et al., 2006). We reanalyzed a scRNAseq dataset of developing mouse periaortic BAT and observed a population of Gata6-positive BA precursors closely related to the one identified in the interscapular BAT. Interestingly, perioaortic BAT receives little contribution from Pax3+ cells and has no contribution from Myf5+ cells (Sanchez-Gurmaches and Guertin, 2014). This therefore suggests that different embryonic populations can contribute to the Gata6-positive brown adipocyte precursors.

Gata6 (GATA binding protein 6), is a zinc finger transcription factor which plays an essential role in the development of extraembryonic tissues (Koutsourakis et al., 1999). Its null mutation in mouse leads to lethality shortly after implantation. Gata6 has also been implicated in the development of several organs including heart, lung, gut, pancreas and skin but its expression during brown fat development has so far not been reported (Freyer et al., 2015; Morrisey et al., 1996). Gata6 was however identified in silico as a transcription factor potentially regulating the expression of brown fat specific genes such as Zic1 and the TCA cycle enzymes such as citrate synthase (Cs) in the ProFat database (Cheng et al., 2018). Also, a GATA recognition motif was identified in ATACseq data of differentiating human adipocytes in vitro (Zhang et al., 2020). In cardiomyocytes, GATA-6 was shown to associate with Ppara to regulate the expression of the glucose transporter Glut4 (Yao et al., 2012). Together, these data suggest that Gata6 could play a role in the regulation of energy metabolism in differentiating brown adipocytes.

Our transcriptome analysis of the brown adipocyte lineage development in mouse identified signaling cues and the appropriate timing of administration which allowed us to modify our paraxial mesoderm development protocol to generate brown fat from hPSCs. This also provided us with a benchmark to which the differentiating hPSCs could be compared. Our comparison of the differentiated human adipocytes generated in vitro in our protocol with those generated by two other recently published protocols (Carobbio et al., 2021; Zhang et al., 2020) suggest that the transcriptional signature of the adipocytes generated in our conditions is closer to endogenous fetal brown adipocytes. In humans, BAT activity shows an inverse correlation with body mass index and percentage of total body fat (Cypess et al., 2009; Saito et al., 2009). Moreover, the graft of BAT in mouse leads to improvement of glucose metabolism in mice exposed to high fat diet or made diabetic following treatment with streptozotocin (Gunawardana and Piston, 2012; Stanford et al., 2013). Our work could therefore help the development of cell therapy strategies involving the graft of human brown adipocytes generated in vitro for the treatment of obesity, metabolic syndrome and diabetes.

## AUTHOR CONTRIBUTIONS

JR designed and performed the experiments, and analyzed the data. JC pioneered the in vitro differentiation in mouse embryonic stem cells. FM analyzed the bulk RNA sequencing data under the supervision of BH. CHW performed the heat generation assay under the supervision of YHT. ZAT generated the PAX3-Venus human pluripotent stem cell reporter line. SG provided technical support to JR for in vitro differentiations. AM performed the electron microscopy experiments. IG provided the human fetal tissues. GK provided the Pax7-Cre:Rosa26-mTmG embryos. FW provided the GATA6-CreERT2:Rosa26-tdTomato embryos. JR and OP wrote the manuscript. OP supervised the project.

## ACKNOWLEDGEMENTS

We thank the Pourquié lab members for helpful discussions. We also acknowledge Sam Wolock and Dan Wagner in the Allon Klein lab (Harvard Medical School) for help with the analysis of single cell RNA sequencing data. We thank Anne Loyens for help with electron microscopy and Matteo Battilocchi for help with the Gata6 lineage tracing experiments in mouse. We thank the NeuroTechnology Studio at Brigham and Women’s Hospital for providing Agilent Seahorse XF96 extracellular flux analyzer and InCell instrument access and consultation on data acquisition and data analysis.

## DATA AVAILABILITY

Raw and processed single cell and bulk RNA sequencing data generated and analyzed in this study have been deposited in NCBI GEO under accession number GSE185623 and GSE185518.

Supplementary table 1 – List of marker genes in Leiden clusters identified in E13.5-E15.5 mouse single cell sequencing data (BA cluster used as reference). cluster_n =cluster name, cluster_s=cluster z-score, cluster_l=cluster log fold change, cluster_p=cluster adjusted p value (Wilcoxon-Rank-Sum test, Benjamini-Hochberg correction method).

## METHODS

### Brown adipocyte differentiation

#### Human induced pluripotent stem cell maintenance

NCRM1 (NIH CRM control iPSC line (male)) human iPSC and other cell lines were routinely cultured on Matrigel (Corning, 354263) coated culture plates (Corning, 353046) in mTeSR (Stemcell Technologies, 85850). Upon confluency, cultures were dissociated into single cells using Accutase (Corning, 25-058-C). Cells were seeded on Matrigel coated culture plates in mTeSR supplemented with 10µM Y-27632 dihydrochloride (R&D Systems, 1254/10) at a density of 56,000/cm2 for routine maintenance of iPSC lines. On the following day, medium was replaced with mTeSR only and medium was changed every day. Cultures became confluent every fourth day and were passaged as described above. For freezing, cultures were dissociated into single cells with Accutase and frozen in NutriFreez D10 Cryopreservation Medium (Biological Industries, 01-0020-50).

#### Differentiation

iPSCs were differentiated into presomitic mesoderm as described previously (Chal et al., 2016). Briefly, confluent maintenance cultures were dissociated into single cells using Accutase (Corning, 25-058-C) and cells were seeded at a density of 30,000-33,000/cm2 on Matrigel (Corning, 354263) coated culture plates (Corning, 353046) in mTeSR (Stemcell Technologies, 85850) supplemented with 10µM Y-27632 dihydrochloride (R&D Systems, 1254/10). Next day, the cells formed small compact colonies. On day 0 of differentiation, to induce presomitic mesoderm, cells were treated with CL medium [DMEM/F12 GlutaMAX (Thermo Fisher Scientific, 10565042) + 1% Insulin-Transferrin-Selenium (Gibco, 41400045) + 3µM CHIR 99021 (R&D Systems, 4423) + 0.5µM LDN-193189 (Stemgent, 04-0074)]. Cells were treated with CL medium for 3 days and medium was refreshed every day. On day 3 of differentiation, cells were changed to CLF medium [CL medium + 20ng/ml FGF-2 (PeproTech, 450-33)]. Cells were treated with CLF medium for 3 days and medium was changed every day.

To further differentiate presomitic mesoderm cells to dermomyotomal fate, cells were changed to HIFL medium [DMEM high glucose (Thermo Fisher Scientific, 11965-118) + Penicillin/Streptomycin (Life Technologies, 15140122) + 15% KnockOut™ Serum Replacement (Life Technologies, 10828-028) + NEAA (Thermo Fisher Scientific, 11140-050) + 0.01mM 2-Mercaptoethanol (Life Technologies, 21985-023) + 10ng/ml HGF (PeproTech, 315-23) + 2ng/ml IGF-1 (PeproTech, 250-19) + 20ng/ml FGF-2 (PeproTech, 450-33) + 0.5µM LDN-193189 (Stemgent, 04-0074)]. Cells were treated with HIFL for two days and medium was changed every day. To let the PAX3 expressing dermomyotomal cells into myogenic and non-myogenic lineages, cells were changed to HI medium (HIFL medium without FGF-2 and LDN-193189) and medium was changed every day.

On day 16-30 of differentiation, cells were replated to differentiate cells into brown adipocyte lineage. Cultures were dissociated into single cells with 2.5mg/ml Collagenase, (Type IV, Thermo Fisher Scientific, 17104019) and 0.05% Trypsine EDTA (Thermo Fisher Scientific, 25200-056) in PBS (Gibco, 14190) and filtered through a 30 µm (CellTrics, 04-0042-2316) filter and seeded on Matrigel coated plates at a density of 60,000-100,000/cm2 in BCTFT medium [DMEM high glucose (Thermo Fisher Scientific, 11965-118) + Penicillin/Streptomycin (Life Technologies, 15140122) + 5% KnockOut™ Serum Replacement (Life Technologies, 10828-028) + 1% Insulin-Transferrin-Selenium (Gibco, 41400045) + 10 ng/ml FGF-2 (PeproTech, 450-33) + 10ng/ml BMP7 (Thermo Fisher Scientific, PHC9544) + 20nM Porcn Inhibitor II, C59 ( Millipore Sigma, 500496) + 10ng/ml TGFb1 (PeproTech, 100-21-10) + 2nM 3,3′,5-Triiodo-L-thyronine sodium salt (Sigma-Aldrich, T6397)]. Cells were treated with BCTFT for 4-6 days and medium was refreshed every day. Adipocyte precursors were either frozen to be used later or differentiated into adipocytes. For freezing, cultures were dissociated into single cells using 0.05% Trypsine EDTA (Thermo Fisher Scientific, 25200-056) in PBS (Gibco, 14190) and cells were frozen in NutriFreez D10 Cryopreservation Medium (Biological Industries, 01-0020-50).

To differentiate adipocyte precursors into adipocytes, cells were cultured in adipogenic medium [DMEM high glucose (Thermo Fisher Scientific, 11965-118) + Penicillin/Streptomycin (Life Technologies, 15140122) + 5% KnockOut™ Serum Replacement (Life Technologies, 10828-028) + 1% Insulin-Transferrin-Selenium (Gibco, 41400045) + 500μM IBMX (3-Isobutyl-1-methylxanthine, Sigma-Aldrich, I7018) + 25.5μg/ml L-Ascorbic acid (Sigma-Aldrich, A4544) + 2nM T3 (3,3′,5-Triiodo-L-thyronine sodium salt, Sigma-Aldrich, T6397) + 5μM TGFb inhibitor SB431542 (Selleck Chemicals, S1067) + 1μM Dexamethasone (Sigma-Aldrich, D4902) + 10ng/ml EGF (Peprotech, AF-100-15) + 4μg/ml Hydrocortisone (Sigma-Aldrich, H0888) + 1μM Rosiglitazone (Sigma-Aldrich, R2408)] for 30-40 days. Medium was changed every third day.

To perform functional assays and immunofluorescence analysis, 30-40 day old replated cultures were dissociated with 2.5mg/ml Collagenase, (Type IV, Thermo Fisher Scientific, 17104019) and 0.05% Trypsine EDTA (Thermo Fisher Scientific, 25200-056) in 1X Phosphate Buffered Saline (Gibco, 14190) and cells were seeded onto suitable plate format required for the assay.

### Human fetal tissue

Human fetal tissues were obtained by the University of Washington Birth Defects Research Laboratory (BDRL) under a protocol approved by the University of Washington Institutional Review Board. BAT tissues were isolated at 98 days (H28540), 115 days (H28572), 122 days (H28560), 125 days (H28626) and 135 days estimated post-conceptual age from the interscapular and scapular region. Tissues were rinsed in 1X Hanks’ Balanced Salt Solution (HBSS, Thermo Fisher Scientific, 14185052) before processing. For immunofluorescence analysis, tissues were fixed with 4% Paraformaldehyde (Electron Microscopy Sciences, 15710) overnight at 4°C, washed 3 times in 1X Phosphate Buffered Saline (PBS, Sigma-Aldrich, P5493) and stored at 4°C in PBS supplemented with 0.1% Sodium Azide (Sigma-Aldrich, 71290) until further processing. For bulk mRNA sequencing, tissues were snap frozen in liquid nitrogen and stored at -80°C until further processing.

### Mouse lines

Pax7-iCre/+;Rosa26-mTmG/+ embryos were generated by crossing Rosa26-mTmG/mTmG (Muzumdar et al., 2007) males with Pax7-iCre/+ females in timed matings and embryos were collected at E14.5. Gata6-EGFPCreERT2 mice were generated by knocking an EGFPCreERT2 cassette into the endogenous Gata6 locus as reported previously (Donati et al., 2017). They were crossed with Rosa26-fl/STOP/fl-tdTomato mice (Madisen et al., 2010). For lineage tracing experiments, pregnant females were injected intraperitoneally with 50 lg/g of tamoxifen (Sigma) at E12.5 and embryos were collected at E15.5. For experiments other than lineage tracing, wildtype CD1 IGS mice (Charles River) were used.

### Generation of UCP1-mCherry and PAX3-Venus reporter line

UCP1 gene was targeted using the CRISPR-Cas9 system-based genome editing to generate a reporter NCRM1 iPSC line which already had been modified at PAX7 locus to introduce Venus. To target the UCP1 locus, a single guide RNA (sense = CACCGGGTTTGCTGCCCGGCGGAC, antisense = AAACGTCCGCCGGGCAGCAAACCC) targeting the 5’ region of the gene was designed using the MIT Crispr Design Tool (www.crispr.mit.edu). The guide RNA was cloned into PSpCas9 (BB)-2A-GFP(PX458) (Addgene, 48138) following protocol from Ran et al., 2013. The final vector was sequence to ensure no mutations were generated during cloning. To generate a targeting vector for homology dependent repair, we cloned 5’ and 3’ 1kb long homology sequence (HA) flanking a (nuclear localization sequence) NLS region from H2B gene sequence, fluorescent protein mCherry sequence and self-cleaving P2A peptide sequence (5’HA-H2B-mCherry-P2A-3’HA) in a pUC19 vector backbone using Gibson Assembly (New England Biolabs (NEB), E5510S). To mutate the PAM site for the guide RNA, the assembled targeting vector was mutated using site-directed mutagenesis using In-Fusion HD Cloning Plus (Takara, 638909). NCRM1 iPSCs were transfected with both Guide RNA vector (PSpCas9 (BB)-2A-GFP) and targeting vector (pUC19-5’HA-H2B-mCherry-P2A-3’HA) using Lipofectamine™ Stem Transfection Reagent (Thermo Fisher Scientific, STEM00001). 24-hours after transfection, cells were sorted using flow cytometry (S3 cell sorter, Biorad) for GFP positive cells and plated at low density for clonal expansion in Matrigel (Corning, #354263) coated culture plates (Corning, 353046) in mTeSR (Stemcell Technologies, 85850) supplemented with 10µM Y-27632 dihydrochloride (R&D Systems, 1254/10) and Penicillin/Streptomycin (Life Technologies, 15140122). After appearance of small colonies, the colonies were sub-cultured and genotyped using PCR for targeted homozygous insertion of the H2B-mCherry-P2A in the UCP1 locus before the transcription start site. Positive clones were sequenced and clones with no undesired mutation were further validated using immunofluorescence and RT-qPCR. Two positive clones were differentiated into adipocytes and expression of UCP1 in mitochondria and mCherry in the nucleus was confirmed. To generate the human PAX3-Venus iPSC line using the CRISPR/Cas9 technology we followed a similar strategy. We generated a Venus knock-in allele by inserting the Venus sequence in front of the coding sequence of exon 1. Guide RNA (sense = 5’-CCGGCCAGCGTGGTCATCCT GGG-3’, antisense = 5’-TGCCCCCAGGATGACCACGC TGG-3’) targeting the 5’ region of the gene was designed using the MIT Crispr Design Tool (www.crispr.mit.edu). The targeting plasmid (pBSKS-2A-3xNLS) was designed to contain NLS sequence, 1.5 kb of the 5’ genomic region of the PAX3 gene, 1 kb of the 3’ sequence and 2A sequence. Targeting vector, together with the Cas9 plasmid was electroporated into cells by nucleofection and clones were sub-cultured and genotyped using PCR for targeted homozygous insertion of the NLS-Venus-2A in the PAX3 locus.

### RNA extraction, Reverse transcription and real time quantitative PCR

Samples were harvested and RNA was extracted using NucleoSpin® RNA kit (Macherey and Nagel, 740955) following manufacturer’s instructions. DNA digestion was performed on column and RNA quality and concentration was measured using Nanodrop. For reverse transcription, 1ug RNA was in a 20ul reaction volume to generate cDNA using iScript™ cDNA Synthesis Kit (Bio-Rad, 1708891). Typically, cDNA was diluted 1:10 with nuclease free water. For real time quantitative PCR, 3.5ul of cDNA, 1.5ul of 10μM forward and reverse primer mix (300μM of forward and reverse primers) and 5ul of iTaq™ Universal SYBR® Green Supermix (Bio-Rad, 172-5124) was used for 10ul reaction volume. For each sample and gene primer set, 3 technical replicates were performed. PCR primers were designed using primer 3, typically spanning splice junctions wherever possible. Primers were validated for amplification efficiency and specificity using a standard curve and melting curves, respectively. PCRs were run on a Bio-Rad CFX384 thermocycler with the following cycling program: initial denaturation step (95°C for 1 minute), 40 cycles of amplification and SYBR green signal detection (denaturation at 95°C for 5 seconds, annealing/extension and plate read at 60°C for 40 seconds), followed by final rounds of gradient annealing from 65°C to 95°C to generate dissociation curves. Relative gene expression was calculated using ΔΔCt method and RPL37A (ribosomal protein L37a) was used as a housekeeping gene. A list of qPCR primers is provided.

RT-qPCR primers:

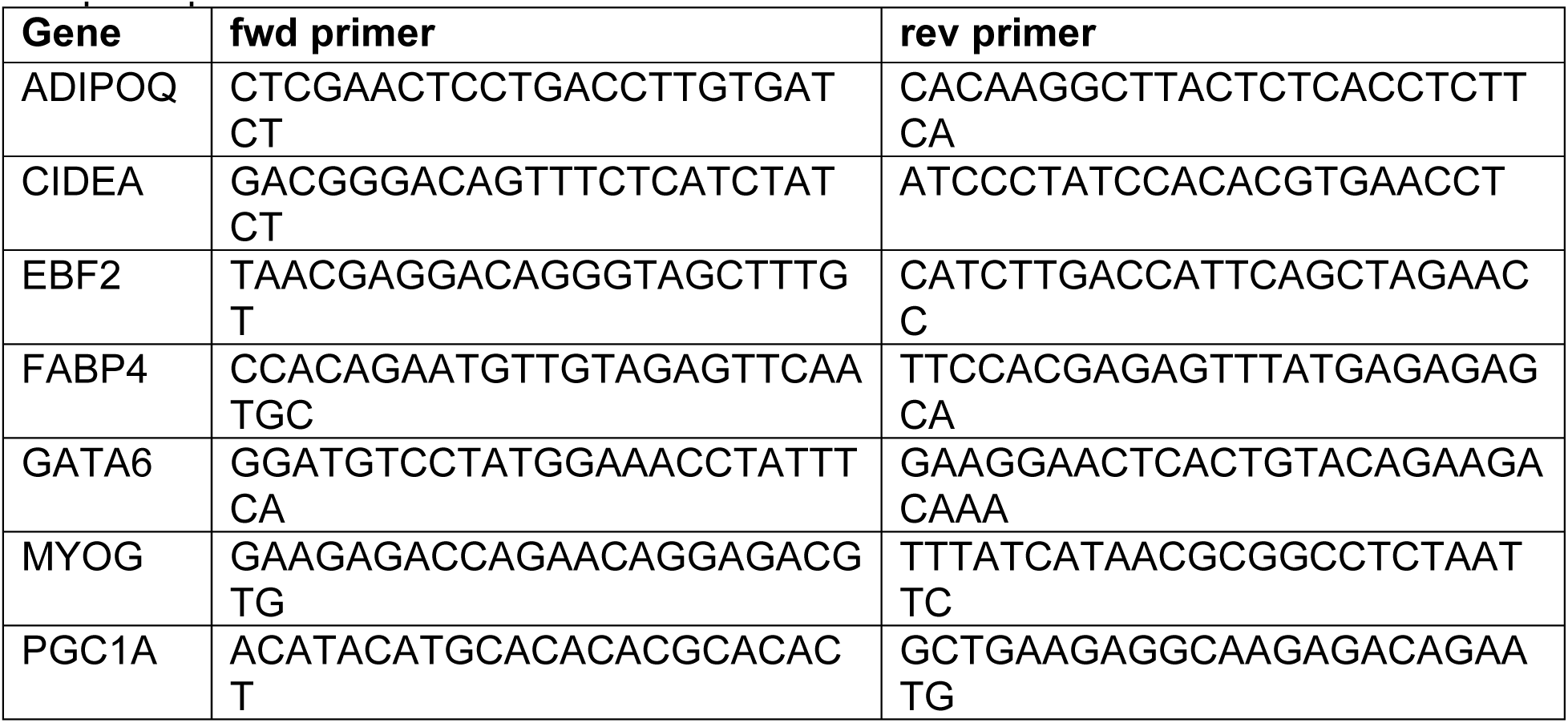
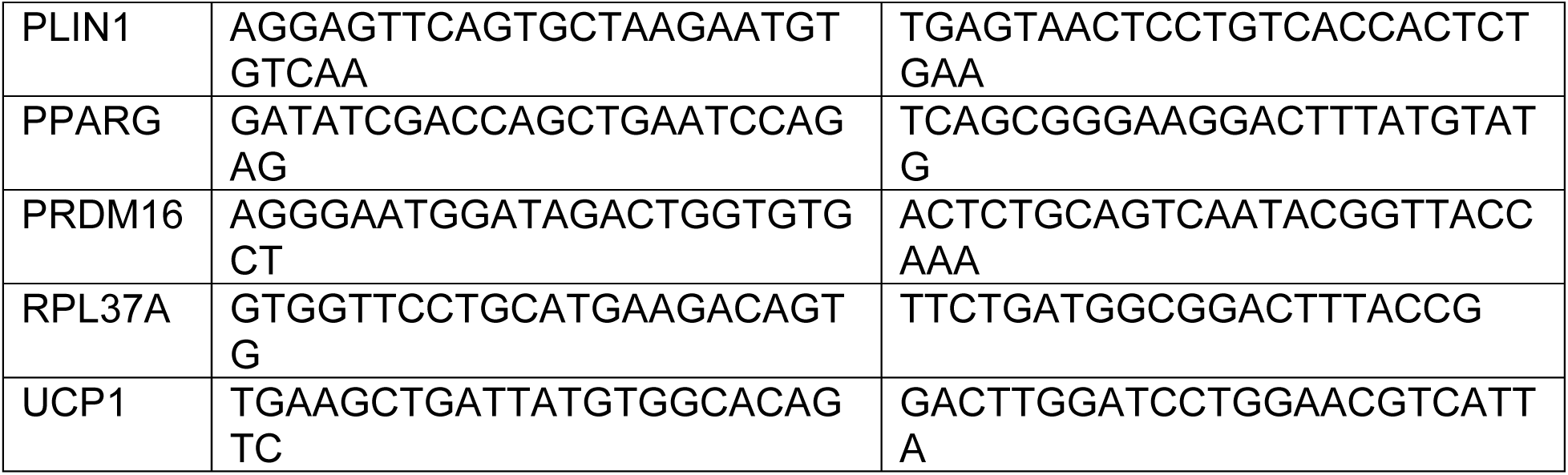

### Immunocytochemistry

#### Cultured cells

Cells were cultured on Matrigel coated plastic culture plates (Corning, #353046), glass bottom 24-well plates (Cellvis, P24-1.5H-N), µ-Dish 35 mm (Ibidi, 81156) or µ-Plate 24 Well Black (Ibidi, 82406). Cultures were washed with 1X Phosphate Buffered Saline (PBS, Sigma-Aldrich, P5493) and fixed with 4% Paraformaldehyde (Electron Microscopy Sciences, 15710) for 20 minutes at room temperature. Cultures were washed with PBS and stored in PBS supplemented with 0.2% Sodium Azide (Sigma-Aldrich, 71290) until used for immunostaining.

#### Mouse tissues

Mouse embryos were dissected out at different stages in Hanks’ Balanced Salt Solution (HBSS) (Gibco, 14185-052). After several washes in HBBS, embryos were fixed in 4% Paraformaldehyde (Electron Microscopy Sciences, 15710) overnight at 4°C. Embryos were then washed in PBS and transferred to 15% sucrose solution (Sigma-Aldrich, 84097) in PBS overnight at 4°C followed by incubation in 30% sucrose solution for overnight at 4°C. Embryos were then embedded in Tissue-Tek O.C.T. Compound (VWR, 25608-930) and stored at -80°C until sectioned. 10-20μm thick sections were cut and stored at -20°C until used for immunostaining.

Fixed cells or sections were permeabilized using 1% Triton-X (Millipore Sigma, T8787) for 10 minutes at room temperature. Samples were washed in PBS and were incubated in blocking solution [PBS supplemented with 3% donkey serum (Jackson ImmunoResearch, 017-000-121) and 0.1% Triton-X] for 1 hour at room temperature. Samples were then incubated with primary antibody diluted in blocking solution at 4°C overnight. Next day, samples were washed 3 times with PBS for 5 minutes. Samples were then incubated with secondary antibody and Hoechst 33342 (ThermoFisher Scientific, H3570) diluted in blocking solution for 1 hour at room temperature followed by 3 washes with PBS for 5 minutes each. Cultured cell samples were stored in PBS until imaged. Mouse tissue slides were mounted using Fluoromount aqueous mounting medium (Sigma-Aldrich, F4680) and were stored at 4°C until analyzed. Samples were imaged using EVOS FL imaging system (Thermo Fisher Scientific) or LSM780 confocal microscope (Zeiss). A list of primary and secondary antibodies is provided.

Primary antibodies:

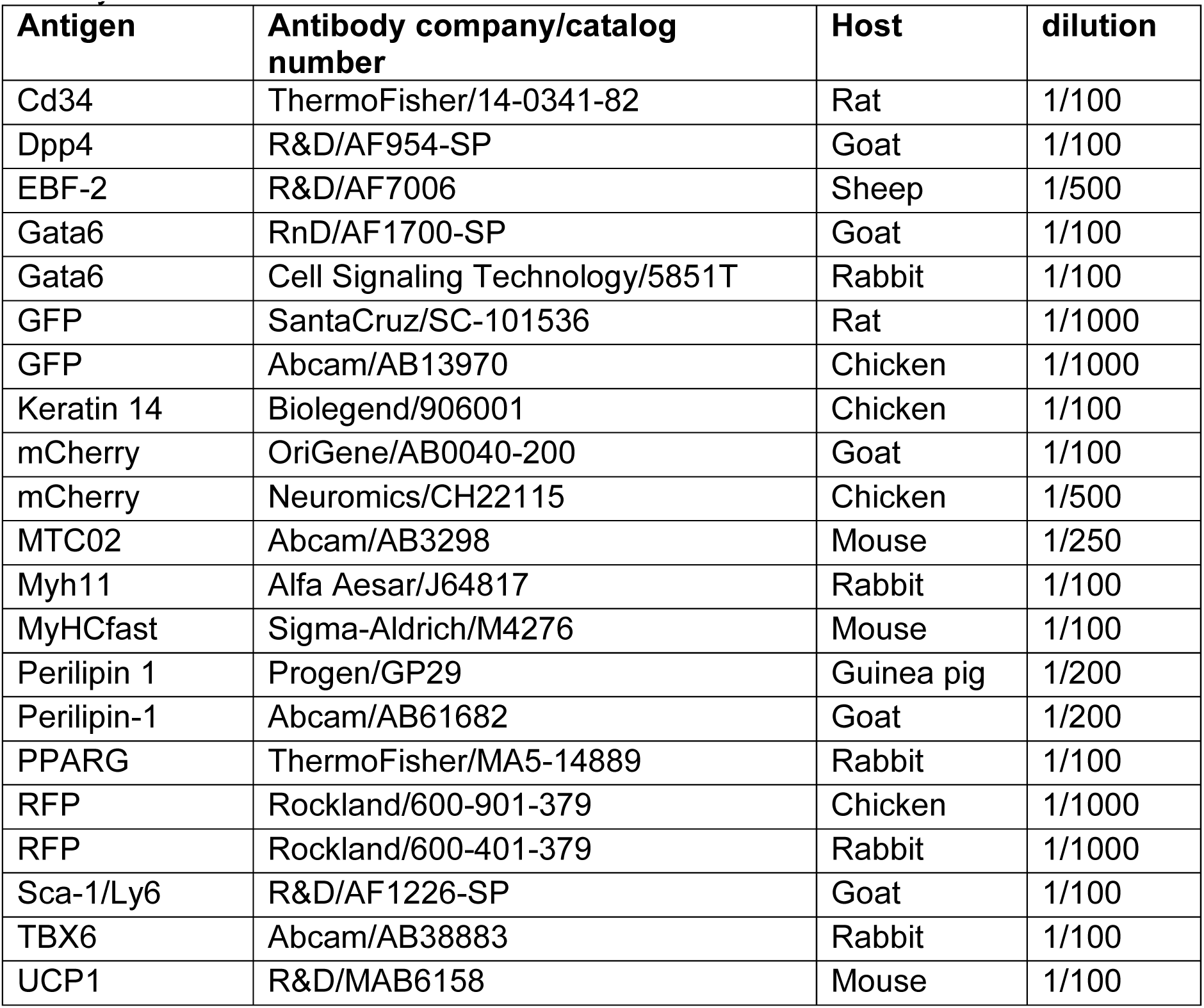

Secondary antibodies:

| **Antibody** | **Company/Catalog number** | **dilution** |
| --- | --- | --- |
| Donkey Anti-Rat IgG H&L (Alexa Fluor® 647) preadsorbed | Abcam/ab150155 | 1/500 |
| Donkey anti-Goat IgG (H+L) Secondary Antibody, Alexa Fluor® 488 conjugate | Thermo Fisher Scientific/A-11055 | 1/500 |
| Donkey anti-Goat IgG (H+L) Cross-Adsorbed Secondary Antibody, Alexa Fluor 647 | Thermo Fisher Scientific/A-21447 | 1/500 |
| Donkey anti-Sheep IgG (H+L) Cross-Adsorbed Secondary Antibody, Alexa Fluor 594 | Thermo Fisher Scientific/A-11016 | 1/500 |
| Donkey anti-Sheep IgG (H+L) Secondary Antibody, Alexa Fluor® 488 conjugate | Life Technologies/A-11015 | 1/500 |
| Donkey anti-Rabbit IgG (H+L) Highly Cross-Adsorbed Secondary Antibody, Alexa Fluor 488 | Thermo Fisher Scientific/A21206 | 1/500 |
| Donkey anti-Rabbit IgG (H+L) Secondary Antibody, Alexa Fluor® 594 | Thermo Fisher Scientific/A-21207 | 1/500 |
| Donkey anti-Rabbit IgG (H+L) Secondary Antibody, Alexa Fluor® 647 | Thermo Fisher Scientific/A-31573 | 1/500 |
| Donkey anti-Rabbit IgG (H+L) Highly Cross-Adsorbed Secondary Antibody, Alexa Fluor 555 | Thermo Fisher Scientific/A-31572 | 1/500 |
| Alexa Fluor® 488 AffiniPure Donkey Anti-Chicken IgY (IgG) (H+L) | Jackson ImmunoResearch Laboratories/703-545-155 | 1/500 |
| Cy™3 AffiniPure Donkey Anti-Chicken IgY (IgG) (H+L) | Jackson ImmunoResearch Laboratories/703-165-155 | 1/500 |
| Donkey anti-Mouse IgG (H+L) Highly Cross-Adsorbed Secondary Antibody, Alexa Fluor 647 | Thermo Fisher Scientific/A-31571 | 1/500 |
| Alexa Fluor® 488 AffiniPure Donkey Anti-Guinea Pig IgG (H+L) | Jackson ImmunoResearch Laboratories/706-545-148 | 1/500 |

### Lipid staining

iPSC-BA cultures were washed with 1X Phosphate Buffered Saline (PBS, Sigma-Aldrich, P5493) and fixed with 4% Paraformaldehyde (Electron Microscopy Sciences, 15710) for 20 minutes at room temperature. Cell were then incubated with 0.5mM BODIPY™ (Invitrogen™, D3922) in PBS for 20 minutes at room temperature. Cells were washed 3 times with PBS and visualized with EVOS FL imaging system (Thermo Fisher Scientific).

### Seahorse assay

Agilent Seahorse XFe96 Analyzer was used to measure oxygen consumption in live iPSC-BAs and precursor cells following manufacturer’s instructions. 30,000 iPSC-BAs and precursor cells were cultured on Matrigel (Corning, 354263) coated Seahorse XF96 cell culture microplates (Agilent Technologies, 101085-004). Assay medium was prepared using Seahorse XF DMEM (Agilent Technologies, 103575-100) supplemented with 1 mM pyruvate (Agilent Technologies, 103578-100), 2 mM glutamine (Agilent Technologies, 103579-100), and 10 mM glucose (Agilent Technologies, 103577-100). Basal respiration was measured 3 times in the assay medium. Cells were then treated with 1.5µM Oligomycin (Tocris, 4110) and 3 measurement were taken. Cells were then treated with 10µM Forskolin (Sigma-Aldrich, F6886) or DMSO (Sigma-Aldrich, D2650) and oxygen consumption rate (OCR) was measured for 70 minutes (12 measurements).

To block the electron transport chain, cells were then treated with 1µM Rotenone (Sigma-Aldrich, R8875) and Antimycin A (Sigma-Aldrich, A8674) for 3 measurements. Cells were then washed with 1X Phosphate Buffered Saline (PBS, Sigma-Aldrich, P5493) and fixed with 4% Paraformaldehyde (Electron Microscopy Sciences, 15710) for 10 minutes at room temperature. Seahorse XF96 cell culture microplates were imaged for DAPI and mCherry using GE INCELL Analyzer 2200 Widefield High-Content Imager. Cells were automatically segmented using custom code in Fiji and number of DAPI and mCherry positive cells were quantified. OCR data from each well was normalized with DAPI positive cell number in each well. Four replicates were performed. Each experiment consisted of 40 technical replicates for each cell type.

### Thermogenesis assay

iPSC-BAs and iPSC precursors were cultured on Matrigel (Corning, 354263) coated 96-well Black Clear-Bottom Plates (Corning, 3603) and were incubated in DMEM/H with 250 nM ERthermAC (Sigma, SCT057) for 30 min at 37°C. After washing with PBS, fresh 90µl DMEM/H without phenol red was added prior to imaging. Fluorescence in stained cells were detected with a GloMax Discover Multimode Detection System (Promega) using 520 nm excitation and emission at 590 nm. Temperature inside the machine was equilibrated at 25°C. After measuring 3 points of basal fluorescence, 10µl forskolin (final concentration 10µM, F6886, Sigma) was added to initiate thermogenesis.10 µl PBS was added for vehicle control group. Fluorescence was recorded every 5 min over 120 min. Results are interpreted as relative intensity (intensity was normalized to basal measurements).

### Electron microscopy

iPSC-BAs were rinsed with 0.1M Phosphate buffer pH 7.5 and fixed in 2.5% glutaraldehyde (Sigma, G7651) in 0.1M phosphate buffer at 4°C, for 2 days. Samples were washed in 0.1M phosphate buffer and post-fixed in 1% osmic acid in 0.1M phosphate buffer for 1 hour at room temperature. Dehydration was performed by serial incubation in 50%, 70%, 80%, 95% and 100% ethanol before incubation in propylene oxide for 30 minutes. Samples were impregnated by incubation in propylene oxide/araldite (1:1 v/v) for 60-90 min, in propylene oxide/araldite (1:2 v/v) for 1 hour and in 100% araldite overnight at 4°C. The samples were finally incubated in Araldite which was allowed to polymerize for 24 hours at 56°C. Ultrathin sections (85nm) were cut with a ultracut Leica EM UC. Sections were contrasted 8 min with 2% uranyl acetate and 8 min with lead citrate (Reynolds). All pictures were taken with the Zeiss EM 900 microscope and a GATAN camera (Orius SC 1000).

### Glycerol release assay

iPSC-BAs were cultured in a (Corning, 354263) coated 96-well culture plate (Genesee Scientific, 25-109). Cells were serum starved for 16 hours and cultured in serum-free DMEM/F12 GlutaMAX (Thermo Fisher Scientific, 10565042) containing 0.5% BSA. Next day, cells were washed with KRB-HEPES buffer [118.5 mM NaCl, 4.75 mM KCl, 1.92 mM CaCl_2_, 1.19 mM KH2PO_4_, 1.19 mM MgSO_4_, 25 mM NaHCO_3_, 6 mM glucose and 10 mM HEPES, pH 7.4] containing 4% fatty-acid-free BSA (Bioworld, 22070017-1). Cells were treated with DMSO or 10µM Forskolin in KRB-HEPES buffer supplemented with 4% fatty-acid-free BSA for at 37°C, 5% CO_2 for_ 4 hours. Cell culture medium was collected for glycerol measurement using the free glycerol reagent (Sigma-Aldrich, F6428) following manufacturer’s instructions. A standard curve was generated using Glycerol Standard Solution (Sigma-Aldrich, G7793). Test samples, standards and water control were incubated with the Free Glycerol Reagent for 5 minutes at 37°C and absorbance was recorded at A540 using a spectrophotometer. To normalize with total protein content, Bradford assay was performed using DC Protein Assay (Biorad, 500-0116) following manufacturers protocol.

### Periodic Acid-Schiff staining

To stain glycogen in differentiating iPSC-BAs, cells were treated with Periodic Acid-Schiff stain (Sigma, 395B). iPSC-BAs were fixed with 4% paraformaldehyde (Electron Microscopy Sciences, 15710) at room temperature for 15 minutes. Fixed cells were washed several times with distilled water and immersed in Periodic Acid Solution for 5 minutes at room temperature. Cells were then washed in distilled water and incubated with Schiff’s Reagent for 15 minutes at room temperature. Cells were washed in running tap water for 5 minutes. Stained cells were visualized in brightfield using EVOS FL imaging system (Thermo Fisher Scientific).

### Bulk mRNA sequencing

UCP1-mCherry knock-in line was differentiated into brown adipocytes as described above. Cells were harvested on day +40 of differentiation and RNA was extracted using NucleoSpin® RNA kit (Macherey and Nagel, 740955) following manufacturer’s instructions. DNA digestion was performed on column and RNA quality and concentration was measured using Nanodrop. Human fetal tissues were harvested as described above and RNA was extracted using NucleoSpin® RNA kit following manufacturer’s instructions.

RNA library preparations and sequencing reactions were conducted at GENEWIZ, LLC. (South Plainfield, NJ, USA). RNA samples received were quantified using Qubit 2.0 Fluorometer (Life Technologies, Carlsbad, CA, USA) and RNA integrity was checked using Agilent TapeStation 4200 (Agilent Technologies, Palo Alto, CA, USA). RNA sequencing libraries were prepared using the NEBNext Ultra II RNA Library Prep Kit for Illumina using manufacturer’s instructions (NEB, Ipswich, MA, USA). Briefly, mRNAs were initially enriched with Oligod(T) beads. Enriched mRNAs were fragmented for 15 minutes at 94°C. First strand and second strand cDNA were subsequently synthesized. cDNA fragments were end repaired and adenylated at 3’ends, and universal adapters were ligated to cDNA fragments, followed by index addition and library enrichment by PCR with limited cycles. The sequencing library was validated on the Agilent TapeStation (Agilent Technologies, Palo Alto, CA, USA), and quantified by using Qubit 2.0 Fluorometer (Invitrogen, Carlsbad, CA) as well as by quantitative PCR (KAPA Biosystems, Wilmington, MA, USA).

The sequencing libraries were clustered on a single lane of a flowcell. After clustering, the flowcell was loaded on the Illumina HiSeq instrument (4000 or equivalent) according to manufacturer’s instructions. The samples were sequenced using a 2×150bp Paired End (PE) configuration. Image analysis and base calling were conducted by the HiSeq Control Software (HCS). Raw sequence data (.bcl files) generated from Illumina HiSeq was converted into fastq files and de-multiplexed using Illumina’s bcl2fastq 2.17 software. One mismatch was allowed for index sequence identification.

For bulk RNA-seq data analysis, we performed quality control on the sequence data (FastQ files) using FastQC (v0.11.9) and used STAR RNA-seq aligner (v2.7.3a) (Dobin et al., 2013) to map the sequenced reads to the human reference genome (GRCh38 release 101 from ENSEMBL). Mapped reads were quantified using featureCounts (v2.0.1) (Liao et al., 2014). Starting from the raw gene counts, normalization and differential expression analysis were done in the R environment (v3.6.0), using the DESeq2 package (v 1.22.2) (Love et al., 2014). Genes were defined as differentially expressed when the false discovery rate was lower than 0.05. Gene Ontology (GO) enrichment analysis was performed on the differentially expressed genes using EnrichR (Kuleshov et al., 2016), choosing the WikiPathway 2019 Human resource.

To quantify the thermogenic potential of each sample we relied on the ProFAT webtool (Cheng et al., 2018). Starting from the raw count matrix, barplot and heatmap, representing the adipose browning capacity of each sample, were generated as standard output of ProFAT.

To produce the heat-map for genes of interest (linked to muscle, pluripotency and brown adipose tissue) we used normalized read counts produced by DESeq2. To calculate up- or down-regulation, we computed the log difference of the average of the biological replicates against the baseline value (average over all conditions) for each gene. All data generated in this study are available at GSEXXXX.

### Single cell RNA sequencing

#### Preparation of single cell suspension

Single cell analysis of differentiating iPSCs and mouse embryos cells was performed using inDrops as previously reported (Klein et al.).

UCP1-mCherry iPSC line was differentiated into brown adipocytes as described above. Cultures were harvested on day 6, 10, 15, 20, 30 of the primary differentiation and day 20 and 40 after replating.

The cultures were washed with PBS (Gibco, 14190) and dissociated into single cells using 2.5mg/ml Collagenase, (Type IV, Thermo Fisher Scientific, 17104019) and 0.05% Trypsine EDTA (Thermo Fisher Scientific, 25200-056) in PBS for 5-15 minutes at 37°C. Dissociated cells were run through 70µm cell strainer (Celltreat, 229483) followed by 30µm cell strainer (CellTrics, 04-0042-2316), spun at 300g for 5 minutes and resuspended in DMEM (Thermo Fisher Scientific, 11965-118) + 5% fetal bovine serum (VWR, 89510-186). Cells were again spun at 300g for 5 minutes and resuspended in DMEM + 5% fetal bovine serum, this was repeated 2-3 times to remove debris. Final resuspension was made in PBS + 0.1% bovine serum albumin (Gibco, 15260-037). Cell density was adjusted to 200,000 cells/ml. For day 6, 10, 15 and 20, 2500 cells were collected from each sample. For samples 30 day, 20 day after replating and 40 day after replating, at least 3000 cells were collected from each sample from two independent differentiations.

For analyzing mouse embryonic tissues, embryos were collected on embryonic day (E) 13.5, 14.5 and 15.5 from wildtype CD1 IGS mice (Charles River). For each stage, back tissue dorsal to rib cage at the level of forelimb was dissected out. Neural tube/spinal cord and dorsal root ganglion were removed wherever possible. For E15.5, epidermis and underlying dermis was removed before dissociating the tissue. Dissected tissues were washed several times in Hanks’ Balanced Salt Solution (HBSS, Thermo Fisher Scientific, 14170112). Tissues were transferred to microfuge tube with 200µl of 2.5mg/ml Collagenase, (Type IV, Thermo Fisher Scientific, 17104019) and 0.05% Trypsine EDTA (Thermo Fisher Scientific, 25200-056) in PBS (Gibco, 14190), chopped into small pieces using scissors and incubated at 37°C for 15 minutes with intermittent shaking. Tissues were mechanically dissociated by triturating several times using wide-bore 1ml pipette. The resulting cell suspension was mixed with HBBS + 10% FBS, filtered through a 30µm cell strainer and spun at 300g for 5 minutes. To remove red blood cells, cell pellet was resuspended and incubated with RBC Lysis Solution (Qiagen, 158902) for 5 minutes at room temperature. Cells were washed with HBSS + 10% FBS for 2-3 times to remove debris. Final suspension was made in PBS + 0.1% BSA. Cell density was adjusted to 200,000/ml. For each stage, cells were collected from two littermate embryos. For E13.5-14.5 embryo, 3000 cells were encapsulated and sequenced from each embryo. For E15.5, 5000 cells were collected and sequenced from each embryo.

#### Encapsulation, sequencing and analysis

Single cells were encapsulated using inDrops technique as reported previously (Klein et al.). Cells were barcoded using v3 sequencing adapters and were sequenced on an Illumina NextSeq 500 using the NextSeq 75 High Output Kits using standard Illumina sequencing primers and 61 cycles for Read1, 14 cycles for Read2, 8 cycles each for IndexRead1 and IndexRead2. Sequence FASTQ files were processed according to indrops.py pipeline (available at github.com/indrops/indrops). Single cell transcriptomes were mapped to mouse (GRCm38/mm10) and human (GRCh38/hg19) reference transcriptomes. Samtools version 1.3.1, rsem version 1.3.0 and Bowtie version 1.2.2 was used with parameter –e 100.

A weighted histogram of transcript counts per cell barcode vs cell barcode abundance was used to identify transcripts originating from abundant cell barcodes. Only transcript counts originating from abundant cell barcodes were included in downstream analysis. Basic filtering parameters were used to exclude cells expressing <500 genes and genes expressed in less than 3 cells. The filtered counts were normalized by total number of counts for each biological sample. Top 1000 variable genes were identified according to Satija 2015. Cell doublets were identified using Scrublet and filtered out (PMID: 30954476). For each cell, fraction of counts due to mitochondrial genes was determined and cells with >0.2 fraction were filtered out. The cell cycle was scored as in Tirosh et al. 2016 (PMID: 27806376). Each cell was given a cell cycle score based on the expression of G2/M and S phase markers. The cells not expressing the markers from G2/M and S phase were identified to be in G0/G1 stage. Source of variation between the libraries and time points were regressed out using bkknn batch correction function (PMID: 31400197). Single cell data were projected into a low dimensional space by principal component analysis (PCA). UMAP (McInnes et al., 2018) was used to embed the neighborhood graph (PCs = 50). Cell clusters were identified using Louvain (community detection based on optimizing modularity) and Leiden graph-clustering method [PMID: 30914743, Blondel et al, 2008]. Top 500 differentially expressed genes were identified by a Wilcoxon rank-sum test by comparing cells of each cluster with cells of all the other clusters. Genes were considered differentially expressed based on fold-change, minimum expression, and adjusted p-value cutoffs, as indicated.

To calculate the RNA velocity, we processed the E14.5 dataset BAM files through the velocyto pipeline to enumerate spliced and unspliced transcripts and obtained loom files (La Manno et al., 2018). We processed the matrix and filtered out cells and genes based on our standard criteria mentioned above. UMAP was used to embed the neighborhood graph (PCs = 50) and cell clusters were identified using Leiden method. RNA velocity analysis was performed on sub-setted somitic cluster using default parameters as described in the scVelo tutorial (Bergen et al., 2020).

Single cell RNA sequencing data from mouse E18 periaortic (pVAT) brown adipose tissue described in Angueira et al. 2021 was processed through the same processing pipeline as described above. Cell state predictions on single cell clusters from pVAT described in Angueira et al. 2021 were made using a kNN-classifier trained on our E13.5-E15.5 mouse single cell clusters (Diaz-Cuadros et al., 2020).

---

**Supplementary Figure 1.**
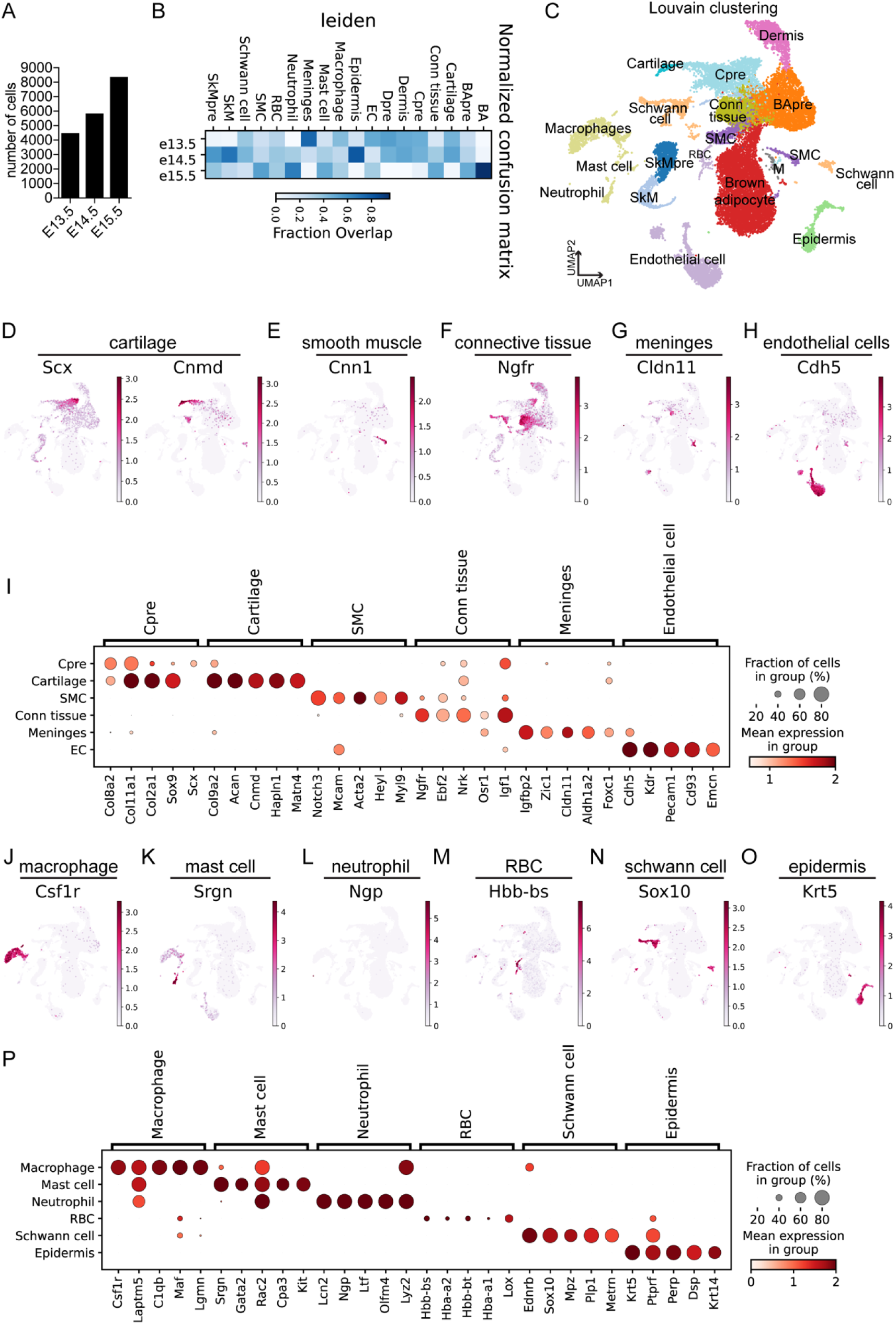
**A** – Bar chart showing number of cells isolated from each stage from developing mouse embryos. The number represents sum of two replicates from each time point. E = embryonic day. **B** – Normalized confusion matrix showing contribution of cells from different time point to identified clusters. **C** – UMAP showing cell clusters identified using Louvain based clustering. Colors indicate identified cell cluster. BApre=brown adipocyte precursors, SkM=skeletal muscle, SkMpre=skeletal muscle precursors, Conn tissue=connective tissue cells, SMC=smooth muscle cells, Cpre=cartilage precursors, M=meninges. **D** – **H** – UMAP plots showing expression pattern of cell cluster specific genes in cartilage (D), smooth muscle cells (E), connective tissue (F), meninges (G), endothelial cells (H). Scale represents log-normalized transcript counts. **I** – Dot plot showing marker genes identified in different clusters (Cpre=Cartilage precursor, SMC=Smooth muscle cells, Conn tissue=connective tissue cells). Scale represents log-normalized transcript counts. **J** – **O** – UMAP plots showing expression pattern of cell cluster specific genes in macrophage (J), mast cell (K), neutrophil (L), RBC, red blood cells (M), Schwann cell (N), epidermis (O). Scale represents log-normalized transcript counts. **P** – Dot plot showing marker genes identified in different clusters (RBC=red blood cells). Scale represents log-normalized transcript counts.

**Supplementary Figure 2.**
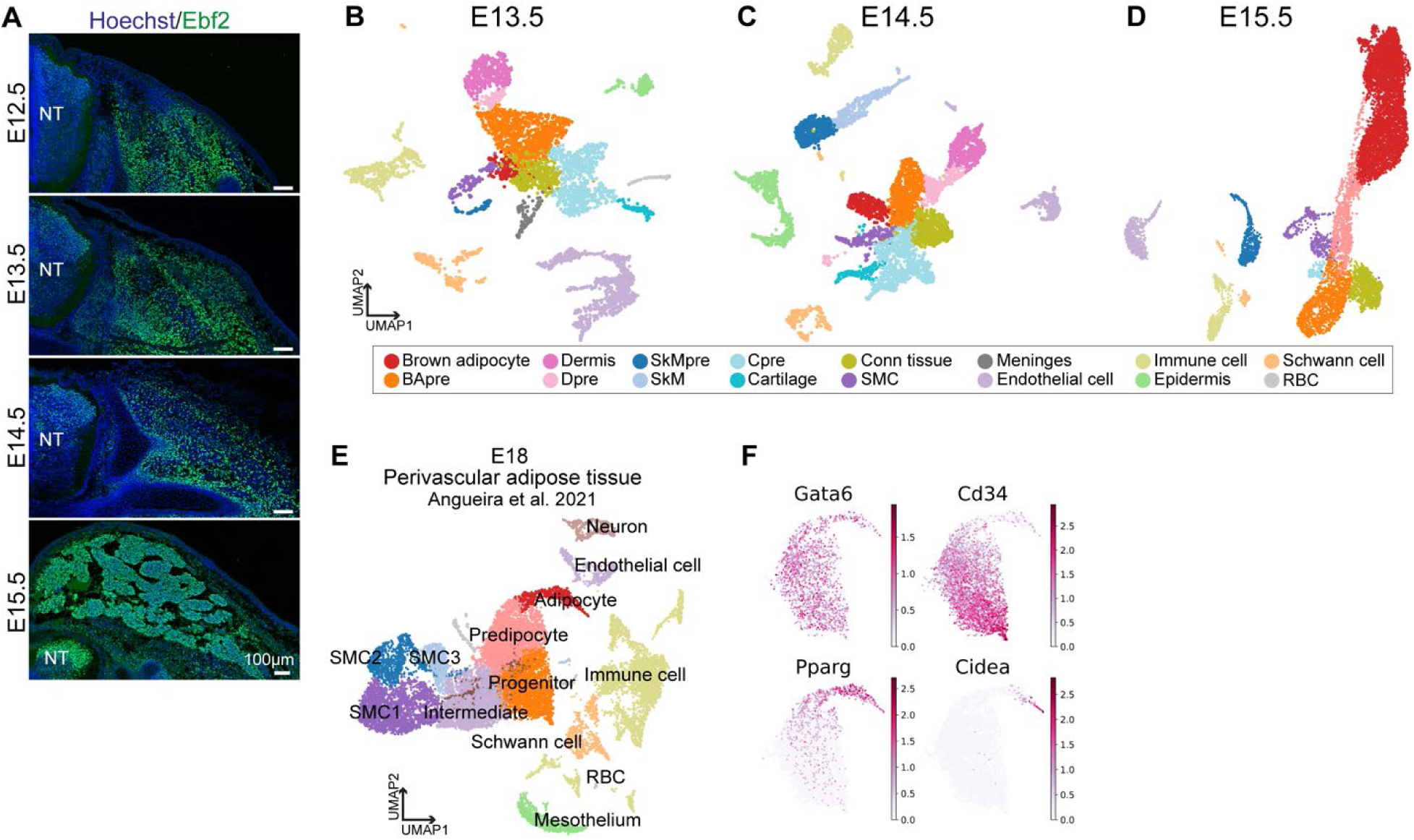
**A** – Immunofluorescence analysis of Ebf2 in developing mouse embryos. Images represent transverse section of mouse embryo at forelimb level during different time points. E=embryonic day, NT=Neural tube. **B** – **D** – UMAP showing cell clusters identified using Leiden based clustering in single cell transcriptomes at E13.5 (50 PC dimensions, 4546 cells) (B), E14.5 (50 PC dimensions, 5396 cells) (C) and E15.5 (50 PC dimensions, 9603 cells) (D). Colors indicate identified cell cluster. BApre=brown adipocyte precursors, Dpre=dermis precursors, SkMpre=skeletal muscle precursors, SkM=skeletal muscle, Cpre=cartilage precursors, Conn tissue=connective tissue cells, SMC=smooth muscle cells, RBC=red blood cells. **E** – UMAP plot generated from mouse single cell transcriptomics data from the developing perivascular adipose tissue on embryonic day 18 described in Angueira et al. 2021. Cells were clustered using Leiden clustering. **F**– UMAP plot showing gene expression patterns in adipogenic cells sub-setted from the single cell transcriptomics data from the developing perivascular adipose tissue on embryonic day 18 described in Angueira et al. 2021. Scale represents log-normalized transcript counts.

**Supplementary Figure 3.**
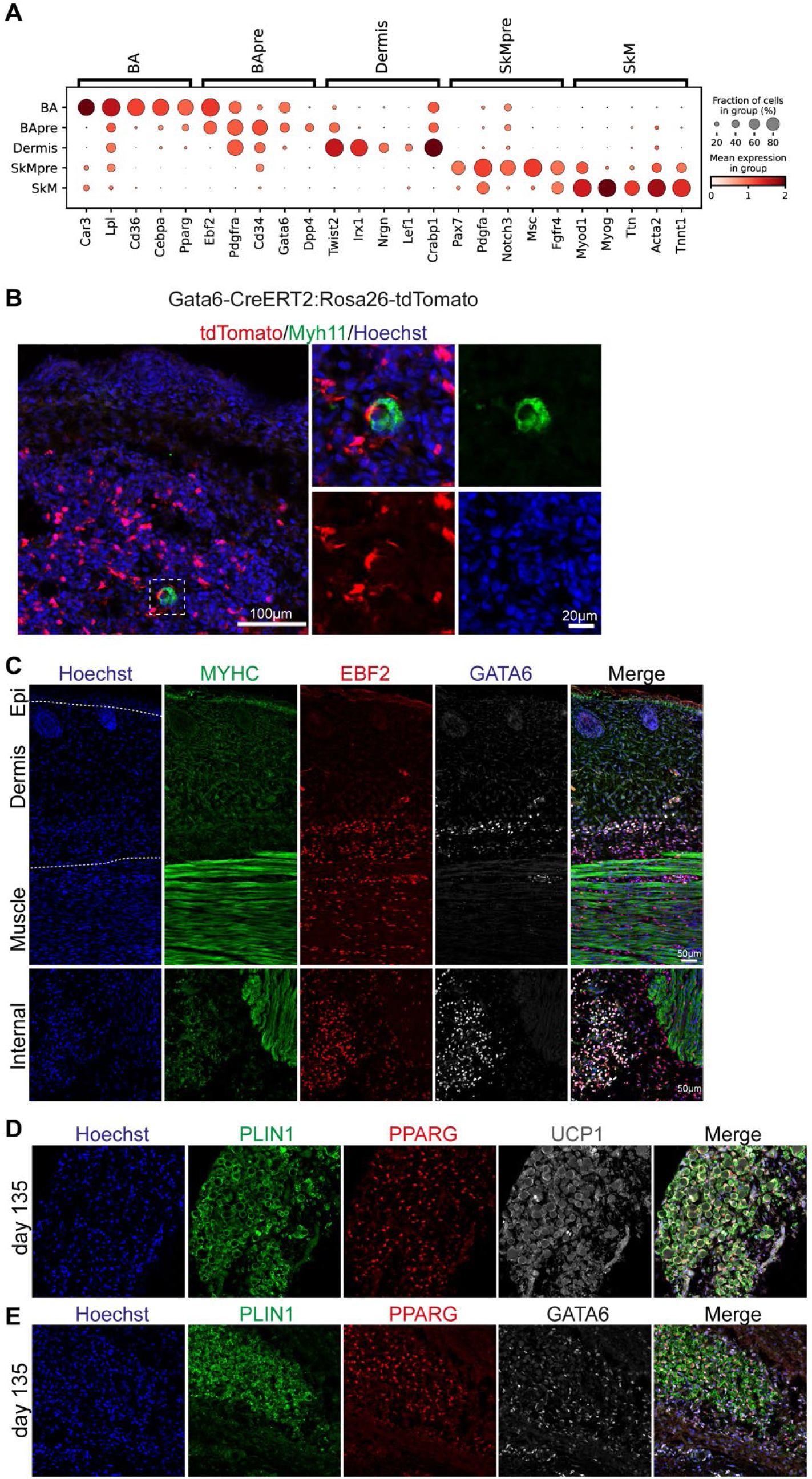
**A** – Dot plot showing marker genes expressed in the brown adipocyte, skeletal muscle and dermis cell clusters at embryonic day 14.5. BA=Brown adipocyte, BApre=Brown adipocyte precursor, SkMpre=Skeletal muscle precursor, SkM=Skeletal muscle. Scale represents log-normalized transcript counts. **B** – Transverse section of Gata6-CreERT2:Rosa26-tdTomato E15.5 embryo at the forelimb level stained for anti-RFP antibody to detect tdTomato positive cells and Myh11 to detect smooth muscle cells. Inset shows magnified image. **C** – Immunofluorescence analysis of human fetal tissues (upper panel – skin, lower panel - dorsal muscles) isolated from scapular region of a 98-day old fetus. Antibody staining for MYHC showing skeletal muscle bundles just below the dermis and internal sections. EBF2 and GATA6 label precursor cells. **D** – **E** – Immunofluorescence analysis of human fetal brown adipose tissue isolated from scapular region of a 135-day old fetus. Expression of PLIN1, PPARG, UCP1 (C), and GATA6 (D) detected with specific antibodies.

**Supplementary Figure 4.**
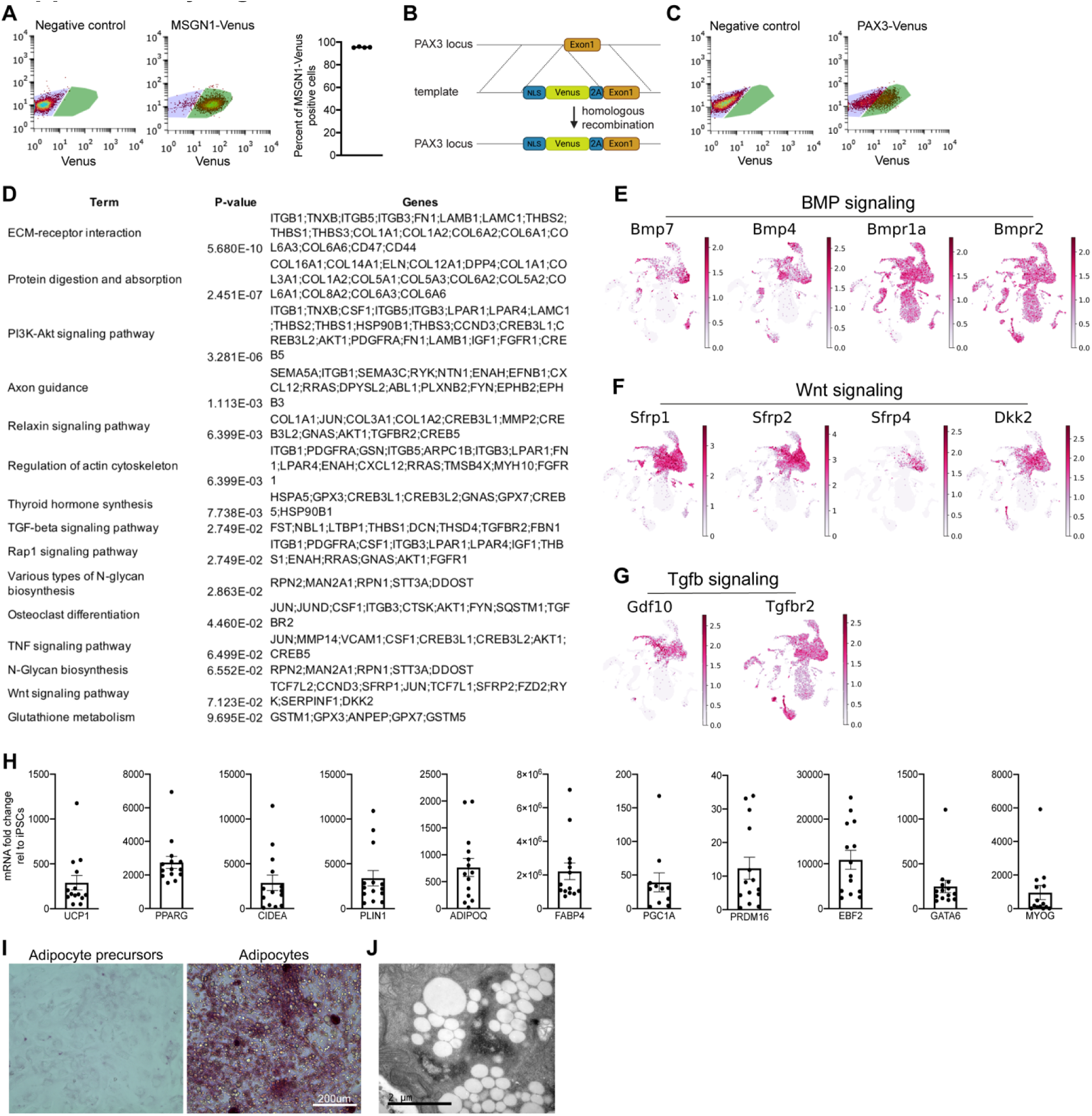
**A** – Flow cytometry analysis of day 2 of differentiating iPSC cultures to measure fraction of Mesogenin-Venus positive cells. Mean ± SD, n=4. **B** – Schematic illustrating gene targeting strategy to generate PAX3-Venus knock-in iPSC line. **C** – Representative flow cytometry plots showing gating strategy for quantifying PAX3-Venus positive cells on day 8 of differentiation. **D** – List of KEGG signaling pathway analysis using EnrichR. Differentially expressed genes in BApre cluster in mouse transcriptomics data were used as an input. **E** – **G** – UMAP plots showing expression patterns of genes involved in BMP (E), Wnt (F) and TGFβ (G) signaling in mouse temporal single cell data. Scale represents log-normalized transcript counts. **H** – RT-qPCR analysis of iPSC-derived brown adipocyte cultures on day 40 after replating. Mean ± SD, n=11-14. **I** – Periodic Acid staining on brown adipocyte precursors (day 20) and brown adipocytes (day 40 after replating). **J** – Representative transmission electron micrographs demonstrating glycogen accumulation in 40 day old replated iPSC-derived adipocytes.

**Supplementary Figure 5.**
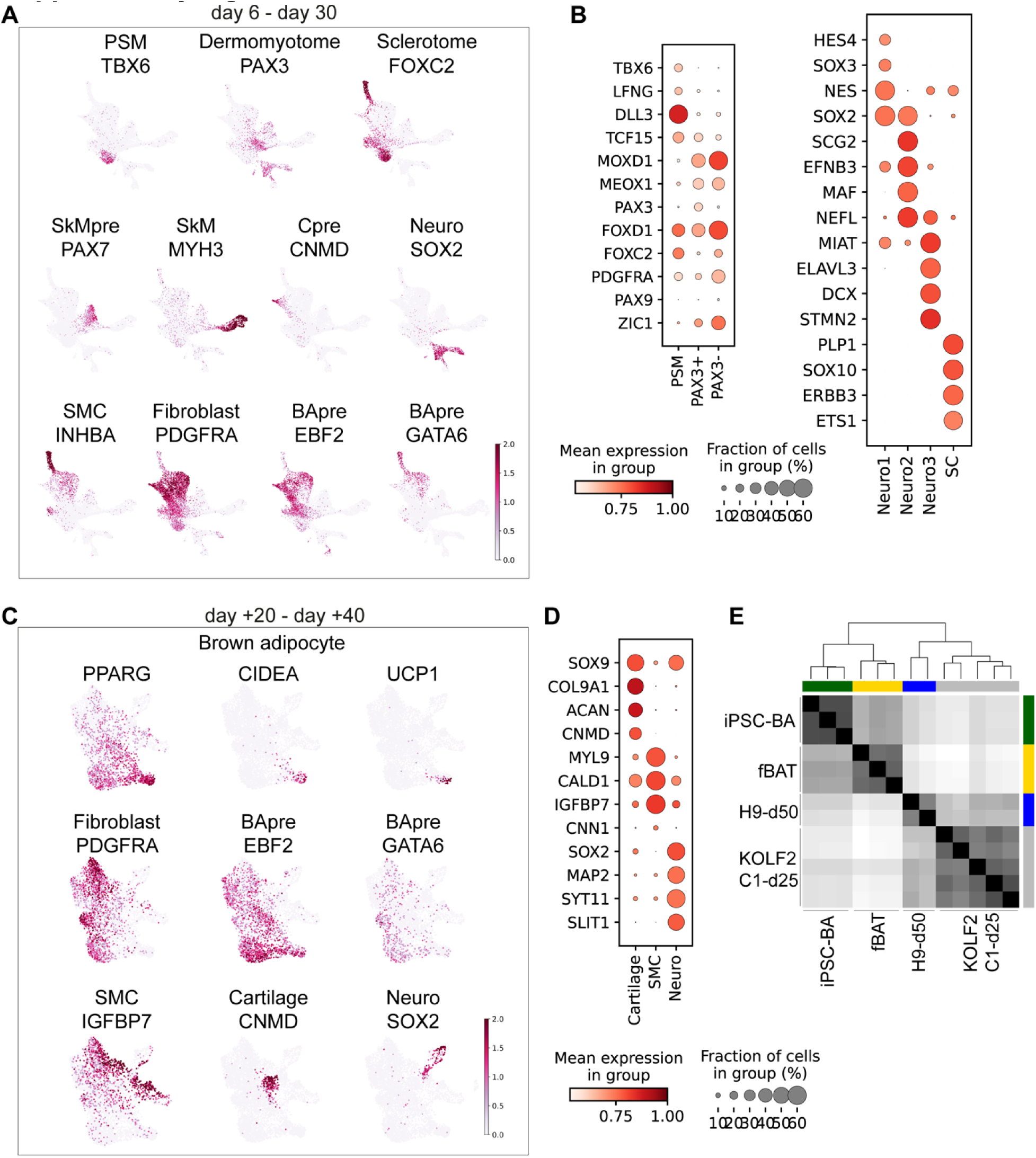
**A** – **B** – UMAP plots and dot plot showing expression pattern of genes in cell clusters identified in the single cell transcriptomes of in vitro differentiated iPSCs from day 6 to day 30. Scale represents log-normalized transcript counts. PSM=anterior presomitic mesoderm, SkMpre=Skeletal muscle precursor, SkM=Skeletal muscle, Cpre=cartilage precursor, Neuro=neural cells, SMC=Smooth muscle cells, BApre= brown adipocyte precursor, SC=Schwann cell. **C** – **D** – UMAP plots and dotplot showing expression pattern of genes in cell clusters identified in the single cell transcriptomes of iPSC derived adipogenic cultures after 20 and 40 days of replating (+ represents days after replating). Scale represents log-normalized transcript counts. BApre= brown adipocyte, SkM=Skeletal muscle, Neuro=neural cells, SMC=Smooth muscle cells. **E** – Matrix plot showing transcriptional overlap between brown adipose specific genes in iPSC derived brown adipocytes (iPSC-BA), human fetal brown adipose tissue (fBAT), human embryonic stem cell derived brown adipocytes in Zhang et al (H9-d50) and iPSC derived brown adipocytes in Carobbio et al (KOLF2-C1-d25).

